# PI(4,5)P_2_-dependence of GABA_A_ receptor channel function revealed by optogenetic manipulation of a binding site

**DOI:** 10.64898/2026.04.06.715776

**Authors:** Risa Mori-Kreiner, Rizki Tsari Andriani, Tobias Strasdeit, Junxian Zhou, Naoyuki Miyashita, Yasushige Yonezawa, Takafumi Kawai, Nikolaj Klöcker, Yasushi Okamura

## Abstract

Ionotropic GABA_A_ receptors (GABA_A_Rs) mediate fast inhibitory neurotransmission in mammalian brains. While recent structural studies have identified that phosphatidylinositol 4,5-bisphosphate [PI(4,5)P_2_], a well-established regulator of numerous ion channels, binds to the α1 subunits of GABA_A_Rs, the functional relevance of this binding has remained elusive. Here, we combine electrophysiology, molecular dynamics simulations, and a recently developed caged lysine technology to define the role of PI(4,5)P_2_ in GABA_A_Rs. We show that GABA_A_Rs are insensitive to acute PI(4,5)P_2_ depletions by voltage-sensing phosphatase, but sensitivity is conferred by neutralizing the K311 binding site, indicating high-affinity binding. Caging of K311 by use of genetic code expansion recapitulated phenotypes of K311 mutant, conferring sensitivity to PI(4,5)P_2_ depletion, whereas uncaging restored insensitivity. Furthermore, caging K311 revealed decelerated activation, which then can be accelerated by uncaging. Additionally, PI(4,5)P_2_-dependence extends to glycine receptors, suggesting PI(4,5)P_2_ is an important endogenous phospholipid modulator of inhibitory receptor channels.

## INTRODUCTION

Pentameric ligand-gated ion channels (pLGICs) mediate fast synaptic transmission that modulates neuronal excitability. The pLGIC superfamily includes nicotinic acetylcholine receptors (nAChR), 5-HT_3_ receptors, GABA_A_ receptors (GABA_A_Rs) and glycine receptors (GlyRs), as well as prokaryotic homologues such as GLIC and ELIC. Lipid modulation of pLGICs has long been recognized, with early studies on *Torpedo* nAChRs demonstrating that defined membrane lipids are essential for reconstituting functional channels ^1–5^. Since then, lipid-protein interactions have been observed across the pLGIC superfamily, highlighting their importance in channel stability, gating, and pharmacology ^6^. Among membrane lipids, anionic phosphatidylinositol 4,5-bisphosphate [PI(4,5)P_2_] is the most abundant phosphoinositide residing in the inner leaflet of the plasma membrane, where it serves as a key modulator of many ion channels ^7–9^.

Recent structural studies revealed that PI(4,5)P_2_ binds directly to the α1 subunit of heterologously expressed full-length α1β3γ2L GABA_A_Rs, with the negatively charged headgroup coordinated by basic residues located at the intracellular ends of the α1 subunits ^10^. PI(4,5)P_2_ binding has since been observed in native mouse and human receptors ^11,12^. Despite the structural evidence, functional studies to date have reported that depletion of membrane PI(4,5)P_2_ produces little or no effects on GABA-evoked currents (I_GABA_), leading to the conclusion that PI(4,5)P_2_ influences processes such as receptor trafficking rather than directly modulating channel gating^10,13^.

In this study, we examined the roles of PI(4,5)P_2_ in channel functions of GABA_A_Rs expressed in *Xenopus* oocytes by combining a voltage-sensing phosphatase (Ci-VSP) ^14,15^, a recently established genetic tool that enables optical control of phosphoinositide binding affinity with a photocageable unnatural amino acid (uAA) ^16^, and atomistic molecular dynamics (MD) simulations. We uncovered that the lysine residue of the α1 subunit, K311, plays a pivotal role in the exceptionally high binding affinity for PI(4,5)P_2_ in GABA_A_Rs and this binding is necessary for rapid channel kinetics. Homopentameric α1 GlyRs also exhibit sensitivity to PI(4,5)P_2_ depletion, dependent on a site that corresponds to the K311 of GABA_A_Rs. These findings indicate that PI(4,5)P_2_ serves as a critical component of both GABA_A_R and GlyR channel activity.

## MATERIALS AND METHODS

### Plasmids

Primers containing restriction sites *Spe*I and *Not*I were designed to amplify the coding sequences of GABA_A_R α1, β3, γ2L subunits and GlyR α1 subunit from cDNA prepared from the brain of C57BL/6 mice. For GABA_A_R, the subunit combination of α1β3γ2L was selected because the cryo-EM structure of the full-length human α1β3γ2L GABA_A_R identified PI(4,5)P_2_ binding in the α1 subunit, coordinated by residues located in the M1-M2, post-M3, and pre-M4 loops ^10^. PCR products were cloned into pSD64TF vector via the *Spe*I and *Not*I sites for *Xenopus* oocyte expression. Full-length nucleotide sequences of the cloned subunits were verified by Sanger sequencing and compared with *Mus musculus* reference sequences obtained from the Ensembl genome browser and the NCBI database. The corresponding NCBI accession numbers are: mGABRA1 (NCBI: NP_034380.1), mGABRB3 (NCBI: NP_032097.1), mGABRG2 (NCBI: NP_032099.1), and mGLRAI (NCBI: NP_001277750.1). Wild-type Ci-VSP and the catalytically inactive Ci-VSP C363S mutant in the pSD64TF vector were used as previously described ^15^. Residue numbering used throughout for GABA_A_R α1 subunit (1–429, QPSQDE…APTPHQ; Uniprot P62812) and GlyR α1 subunits (1–422, ARSAPK…EDVHNK; Uniprot Q64018) corresponds to their respective mature polypeptide numbering.

Plasmids used for caged lysine experiments including pE323-HCKRS (encoding the pyrrolysyl-tRNA synthetase and tRNA_CUA_) ^17^ and pCS2^+^-HCKRS (encoding the pyrrolysyl-tRNA synthetase) ^18^ as well as a single strand DNA fragment of pyrrolysyl-tRNA_CUA_ harboring a T7 promoter sequence at the 5’ end ^19^, and the hydroxycoumarin-lysine (HCK) compound ^20^ were used as described previously ^16^.

Site-directed mutagenesis was conducted using PrimeSTAR Max DNA Polymerase with primers designed for each respective mutation (Supplementary Table 2). Mutations were selected based on sequence comparisons among the α subunit isoforms (Fig. 1a). For R248 and K390 (mouse protein numbering for mature α1 subunit), where positive charges are conserved, we substituted glutamine (Q). For K311 and R312, which are not conserved in the α4 and α6 isoforms, we substituted the corresponding residues from the α6 (asparagine, N, and leucine, L, respectively). Construct visualization and primer design were performed using ApE software (A Plasmid Editor)^21^ and In-Fusion Cloning Primer Design Tool (Takara Bio, Japan), respectively. Primers were synthesized by Thermo Fisher Scientific (USA), and all constructs were verified by Sanger sequencing (Genewiz, Azenta Life Sciences, USA).

**Figure 1.**
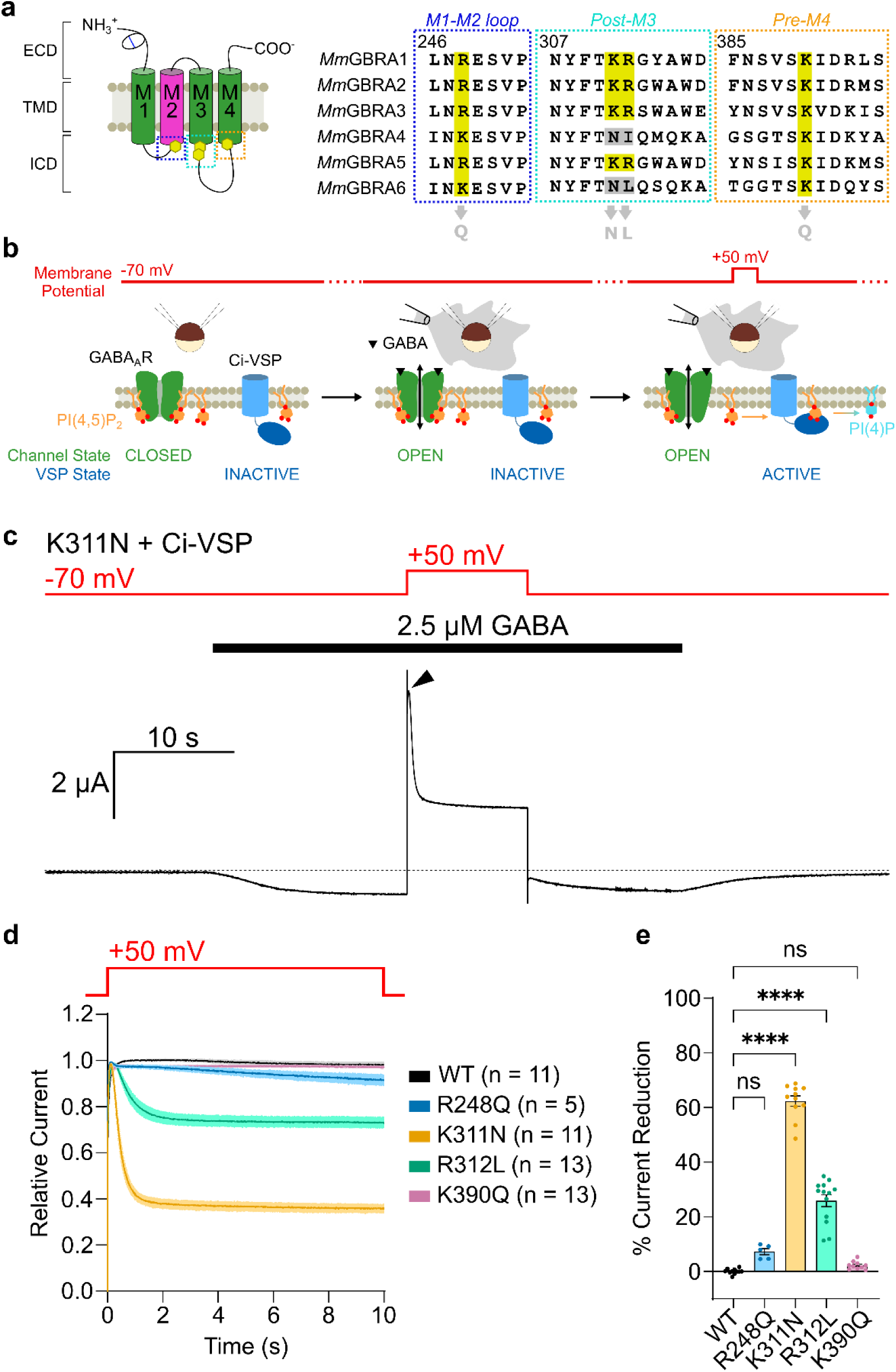
Neutralizing positive charges of PI(4,5)P_2_ binding sites confers sensitivity to PI(4,5)P_2_ depletion. **(a)** GABA_A_R α1 topology (left) with PI(4,5)P_2_ sites (yellow hexagons). Right: α1–α6 alignment; binding sites highlighted (yellow), non-conserved M3 residues and experimental mutations (gray). **(b)** Ci-VSP/TEVC schematic showing membrane potential (red), GABA_A_R state (green), and Ci-VSP state (blue). **(c)** Representative α1K311N GABA_A_R trace; arrow indicates peak outward I_GABA_. **(d)** Normalized outward I_GABA_ time course during 10-s depolarization. Shading: mean ± SEM (n = 5–13 oocytes; 2–3 batches). **(e)** Percent current reduction. Statistical significance was assessed by Kruskal-Wallis test followed by Dunn’s post hoc test (WT vs. α1R248Q, p = 0.128; WT vs. α1K311N, p = <0.0001; WT vs. α1R312L, p = <0.0001; WT vs. α1K390Q, p = 0.599). Data represents mean ± SEM.

For caged lysine experiments, the natural C-terminal *amber* stop codon (TAG) of the GABA_A_R α1 subunit was replaced with an *opal* stop codon (TGA) to avoid unintended incorporation of HCK, generating the GABA_A_R α1*^opal^* construct. For site-specific incorporation of HCK, the desired lysine residues were mutated to TAG codons in the GABA_A_R α1*^opal^* background.

### Oocyte preparation

*Xenopus laevis* were purchased from domestic animal venders, either from Xenopus Youshoku Kyouzai (Ibaraki Prefecture, Japan) or Watanabe Zoushoku (Hyogo Prefecture, Japan). Oocytes were surgically removed from frogs that were under anesthesia of 0.15% ethyl 3-aminobenzoate methane sulfonate (Tokyo Chemical Industry, Japan; CAS RN: 886-86-2). Isolated oocytes were treated with 0.5–2 mg mL^-1^ collagenase P (Roche) rocking for 2–4 hours at 18°C to remove the follicular membrane. After collagenase treatment, stage V-VI oocytes were selected and injected with 50 nL of complementary (cRNA) mixtures using NANOJECT II (Drummond Scientific Company, USA). Alternatively, ready-to-use *Xenopus laevis* oocytes were purchased from EcoCyte Biosciences (Dortmund, Germany) and selected for cRNA injections. Injected oocytes were incubated at 18°C for 2–3 days in ND96 solution containing 96 mM NaCl, 2 mM KCl, 1.8 mM CaCl_2_, 1 mM MgCl_2_, and 5 mM HEPES (pH to 7.4–7.5 with HCl), supplemented with 0.1% gentamycin (Nacalai Tesque, Inc., Japan; CAS RN: 1405-41-0) and 5 mM sodium pyruvate (FUJIFILM Wako, Japan; CAS RN: 113-24-6) prior to electrophysiological recordings.

All experiments were approved by the Animal Care and Use Committee at the University of Osaka Graduate School of Medicine and were performed in accordance with its guidelines.

### cRNA preparation

Constructs in the pSD64TF vector were linearized with *Xba*I and plasmid pCS2^+^-HCKRS was linearized with *Not*I to synthesize cRNAs using mMESSAGE mMACHINE SP6 Transcription Kit (Thermo Fisher Scientific, USA) following manufacturer’s protocol. Synthesis of the pyrrolysyl-tRNA_CUA_ (PylT) from the single DNA fragment containing the T7 promoter sequence was performed according to previous method reported ^18^, using primers (Fw primer: AATACGACTCACTATAGGA; Rv primer: TGGCGGAAACCCCGGGAATCTAA) to amplify the template to use for in vitro transcription using MEGAscript T7 Transcription Kit (Thermo Fisher Scientific, USA) following manufacturer’s protocol ^16^.

For GABA_A_R expression alone, α1, β3, and γ2L cRNA mixture was injected at a ratio of 1:1:10 to ensure the incorporation of the γ2L subunit ^22^. For co-expression with Ci-VSP (wild type or the C363S mutant), cRNAs were mixed at a ratio of 1:1:10:10 (α1:β3:γ2L:Ci-VSP). For GlyR expression, either α1 subunit was injected alone or co-injected with Ci-VSP at a 1:1 ratio.

For experiments combining caged lysine system with the Ci-VSP co-expression technique, we used a Ci-VSP construct that harbors the *ochre* stop codon (TAA) in place of its natural C-terminal TAG stop codon as described previously (Ci-VSP*^ochre^*) ^23^. For co-expression, α1-K311HCK, β3, γ2L, and Ci-VSP*^ochre^* cRNA mixture was injected at a ratio of 20:1:10:6 respectively.

### Unnatural amino acid (uAA) incorporation

Two approaches, nuclear injection or all-RNA injection, were used for the incorporation of HCK into mutant receptors containing TAG stop codons.

For nuclear injection (two-step injection), on day 1, 1.25 ng of plasmid pE323-HCKRS was injected into the nucleus of defolliculated oocytes (50 nL/oocyte). Oocytes were then incubated for 24 h at 18 °C to allow the expression of the pyrrolysyl-tRNA/tRNA_CUA_ system. On day 2, under minimized light exposure, cRNA mixture containing the TAG mutant cRNA was co-injected with 23 nL of 1 mM HCK (dissolved in DMSO). Following injection, oocytes were maintained in dishes wrapped in foil (to ensure dark conditions) and incubated in ND96 solution supplemented with gentamicin and sodium pyruvate for 2–3 days.

For all-RNA injection (the single-step approach), HCKRS cRNA (prepared from the pCS2^+^-HCKRS construct), TAG mutant cRNA, PylT, and the HCK were mixed and co-injected into defolliculated oocytes in a single step. Oocytes were maintained in foil-wrapped dishes under dark conditions and incubated in ND96 supplemented with gentamicin and sodium pyruvate for 2–3 days.

It is worth noting that oocytes injected with either two-step injection or single-step approach to incorporate HCK, resulted in weak membranes compared to oocytes injected with channel RNA only.

### Electrophysiological recordings

Two-electrode voltage-clamp (TEVC) data were acquired with a bath-clamp amplifier (OC-725C, Warner Instruments, USA) controlled by PatchMaster software (HEKA Elektronik, Germany) for pulse protocol manipulation. Output signals of I_GABA_ that were sampled at 1 kHz and VSP sensing currents sampled at 100 kHz, were digitized with an AD/DA converter (LIH 8+8, HEKA Elektronik, Germany). Glass microelectrodes (1.5 outer diameter/1.17 inner diameter, Harvard Apparatus, USA; W3 30-0066) were pulled with a P-97 micropipette puller (Sutter Instruments, USA) and filled with 3 M potassium acetate and 10 mM KCl. Micropipettes used in our experiments had resistances of 0.5–0.8 MΩ.

All TEVC recordings were performed in a high-chloride NMDG solution (100.8 mM NMDG, 5 mM MgCl_2_, and 20 mM HEPES, adjusted to pH 7.4–7.5 with HCl). Oocytes were placed in a small recording chamber and continuously superfused using a gravity-fed perfusion system. Agonist solutions containing GABA (Sigma, USA; CAS RN: 56-12-2) and glycine (FUJIFILM Wako, Japan; CAS RN: 56-40-6) were prepared at varying concentrations by diluting stock solutions (0.05–1 M in water, aliquoted and stored at –20 °C) into the high-chloride NMDG recording solution. Agonists were applied either by gravity-fed perfusion (for experiments that used lower concentrations) or by direct application with an electronic pipette (BM-MPA1200; A&D Company, USA) positioned at the recording chamber for rapid solution exchange. For pipette application, the dispensed volume was set to 5–10 times the chamber volume to ensure complete exchange of the bath solution. To improve the temporal resolution of recordings, we optimized the recording chamber to minimize the bath volume to be approximately 50 µL.

Current responses to agonist applications (GABA or glycine) were recorded from oocytes voltage-clamped at –70 mV in a gap-free protocol. For Ci-VSP co-expression experiments, phosphatase activity was activated by applying a 5–10-s depolarizing pulse to +50 mV, either during or immediately before agonist application. In the experiments of VSP activation, during agonist application, the outward current induced by agonist application was quantified at agonist concentrations not yielding receptor desensitization. Taking the outward current to define the activity of GABA_A_Rs, as compared to taking the inward current, holds the advantage of avoiding contamination by overlapping endogenous currents, which are often evoked inwardly upon repolarization from +50 mV. In the same oocytes, Ci-VSP expression level was verified by measuring the sensing current, which was evoked by depolarizing step pulses from the holding potential of –60 mV to +160 mV in 10-mV increments. P/N leak subtraction was performed using a P/-8 protocol.

For caged lysine experiments, current responses to varying GABA concentrations were recorded from oocytes voltage-clamped at –70 mV. Oocytes with an initial leak current within 30-100 nA were selected to ensure good recording quality given the fragility of the HCK-containing membrane. Uncaging of HCK mutants was achieved by illuminating the recording chamber with UVA-LED lamps. A 365 nm/20 W lamp (SecurityIng) was used for the results shown in Fig. 2c–d and Fig. 3b, whereas a 395 nm/10 W lamp (Alonefire C8) was used for the results shown in Fig. 2e. UV irradiation was monitored with a UV sensor (NF-MTA08UV; OPTEX FA, JAPAN) connected to an output (MDF-TA; OPTEX FA, JAPAN) during recordings. Depending on the protocol, the timing of uncaging varied to assess changes in channel activity between the caged and uncaged states.

**Figure 2.**
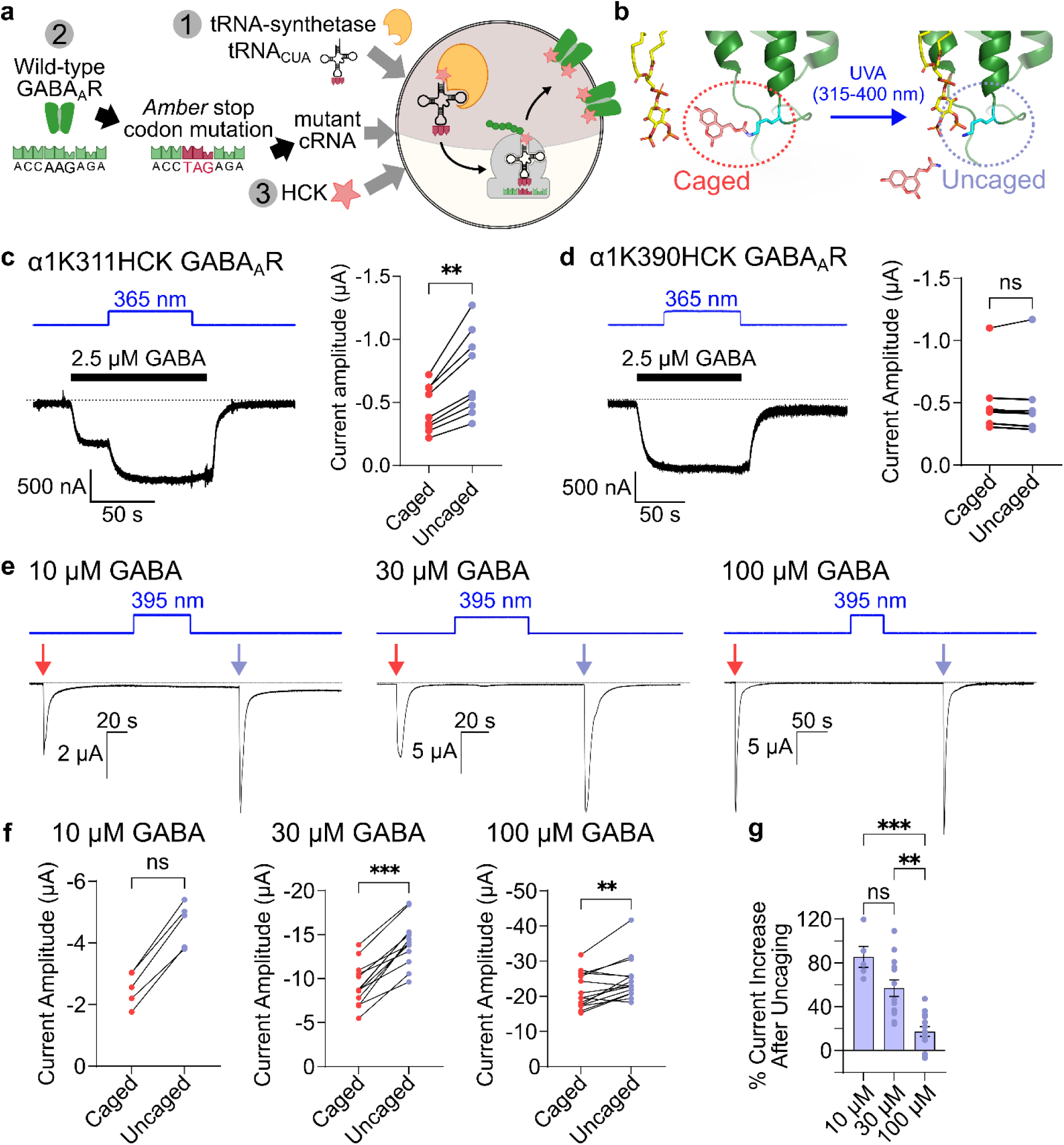
K311 of α1 subunit is a critical residue for GABA_A_R channel function. **(a)** Simplified scheme of HCK incorporation into GABA_A_Rs. **(b)** Structural representation of HCK incorporated at residue position K311. Irradiation by UVA light (315–400 nm wavelength) cleaves the hydroxycoumarin group of HCK, restoring the native lysine side chain at K311. **(c-d)** Representative traces of **(c)** α1K311HCK and **(d)** α1K390HCK uncaging during 2.5 µM GABA application. UV irradiation was performed with a 365 nm/20 W lamp. The absolute current amplitudes before (“Caged”) and after uncaging (“Uncaged”) was plotted and analyzed by Wilcoxon matched-pairs signed rank test (α1K311HCK: p = 0.004; n = 9 oocytes; α1K390HCK: p = 0.375; n = 7; from 3–4 batches). Data represents mean ± SEM. **(e)** Representative traces of α1K311HCK uncaging in a paired pulse protocol. Oocytes were voltage-clamped at –70 mV and exposed to varying GABA concentrations applied twice: before uncaging (pink arrows) and after uncaging (cyan arrows). To uncage, 395 nm/10 W lamp was used. **(f)** Absolute current amplitudes corresponding to **(e)** were plotted for each GABA concentration. Statistical analysis was assessed with Wilcoxon matched-pairs signed rank test (10 µM, p = 0.065; 30 µM, p = 0.0002; 100µM, p = 0.003; n = 5–15). Data represents mean ± SEM. **(g)** Percent current increase after uncaging was calculated for each GABA concentration. Statistical significance was assessed by Kruskal-Wallis test followed by Dunn’s post hoc test (10 µM vs. 30 µM, p = 0.680; 10 µM vs. 100 µM, p = 0.001; 30 µM vs. 100 µM, p = 0.002; n = 5–15). Data represents mean ± SEM.

**Figure 3.**
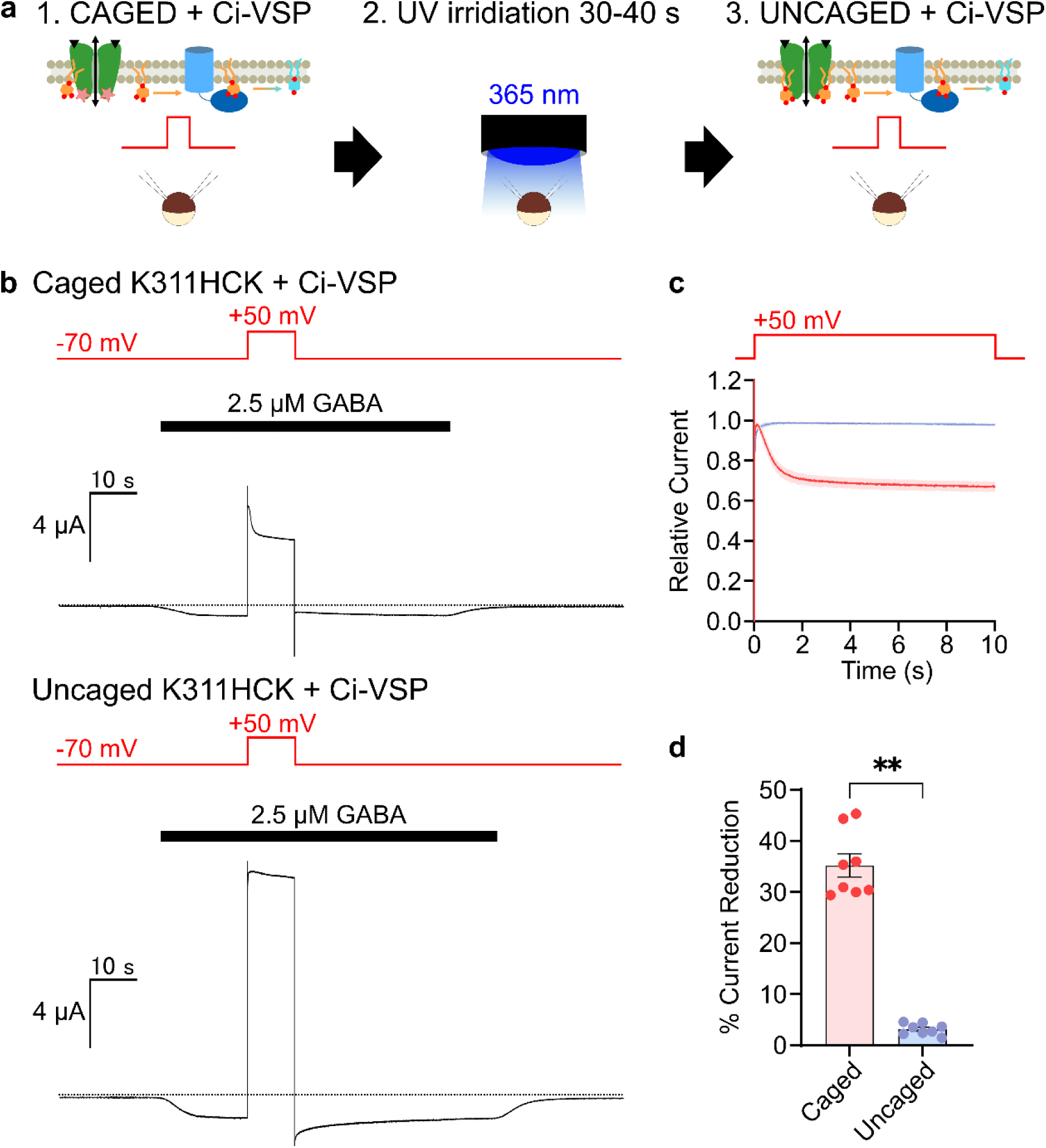
WT GABA_A_Rs require PI(4,5)P_2_ for channel function. **(a)** Schematic overview of caged lysine co-expression with Ci-VSP TEVC experiment. 1) Ci-VSP activation + caged α1K311HCK GABA_A_Rs; 2) uncaging with 365 nm/20 W lamp; 3) Ci-VSP activation + uncaged α1K311HCK GABA_A_Rs. **(b)** Representative trace of Ci-VSP activation in caged α1K311HCK GABA_A_R (top) and uncaged K311HCK GABA_A_R (bottom). **(c)** Time course plot comparing normalized outward I_GABA_ of caged α1K311HCK GABA_A_R (blue) and uncaged α1K311HCK GABA_A_R (pink) during Ci-VSP activation. Plot shows group means ± SEM (shading). **(d)** Comparison of percent current reductions between caged vs uncaged α1K311HCK GABA_A_Rs during Ci-VSP activation. Statistical significance was assessed with Wilcoxon matched-pairs signed rank test (p = 0.008; n = 8). Data represents mean ± SEM.

All current traces were visualized for analysis and figure generation using IgorPro 8 or 9 software (WaveMetrics Inc., USA).

### Outside-out patch clamp recording with rapid perfusion from *Xenopus laevis* oocytes

Patch clamp recordings of *Xenopus* oocytes were performed 1–2 days after injection for WT or 2–3 days after injection for α1K311HCK using an EPC 10 USB patch clamp amplifier operated by Patchmaster software (HEKA, Germany). Rapid extracellular solution exchange was achieved using a piezoelectric linear actuator (PI Physik Instrumente, Karsruhe, Germany) to operate a theta-barrel capillary, with agonist and bath solutions forced through two channels. Solution exchange within 100 µs was confirmed by measuring the 20–80% rise time from open tip responses to changes in external solutions differing in ionic concentrations ^24^. The piezoelectric actuator was connected to an HVPZT amplifier (PI Physik Instrumente, Karlsruhe, Germany) and an LPBF-01GX filter (npi electronic GmbH, Tamm, Germany), controlled by the EPC 10 amplifier and Patchmaster, respectively.

Patch pipettes with resistance ranging from 1–3 MΩ were pulled from borosilicate capillaries and filled with an internal solution containing 140 mM KCl, 10 mM HEPES, 5 mM EGTA, and 2 mM MgCl₂, adjusted to pH 7.4 with KOH. The external solution contained 140 mM NaCl, 10 mM HEPES, and 2 mM MgCl₂, adjusted to pH 7.4 with NaOH, and was filtered through 0.45 μm nylon filters under vacuum. The osmolality of both solutions was adjusted to 285–295 mOsmol/kg using KCl and NaCl or water, respectively. Agonist (1 mM GABA) was dissolved in the external solution.

The vitelline membrane of the oocyte was manually removed using forceps. Recordings were performed at room temperature, and patches were clamped at a holding potential of –70 mV. GABA_A_Rs activation and deactivation were recorded by applying 1 mM GABA for 1 ms with a 2-s interpulse interval, whereas desensitization was recorded by applying 1 mM GABA for 1 s with 10-s interpulse interval. GABA applications were repeated 4–10 times for WT and 2–10 times for α1K311HCK. Due to the fragility of HCK-containing oocyte membranes and limited resources during outside-out patch recordings, data was collected for both 1-ms and 1-s GABA applications from oocytes that remained viable until the final protocol. Current responses were sampled at 50 kHz or 100 kHz using a 2.9 kHz low-pass filter. Uncaging of α1K311HCK GABA_A_Rs was performed by irradiation with 395 nm UV light (10 W, Alonefire C8) for 2 min. The duration of UV light irradiation was determined by balancing uncaging efficiency with cell viability, as prolonged exposure induced phototoxicity. After evaluating various exposure times, a 2-minute duration was found to be optimal for achieving reproducible results. All current traces were visualized for analysis and figure generation using IgorPro 8 or 9 software (WaveMetrics Inc., USA).

### Analysis of GABA- and glycine-evoked outward current during Ci-VSP activation

During membrane depolarization to +50 mV to activate Ci-VSP, agonist-evoked currents in both GABA_A_Rs and GlyRs reverse direction from inward to outward, a process influenced by their intrinsic voltage sensitivity ^25^. To assess the effect of acute PI(4,5)P_2_ depletions by Ci-VSP on channel or transporter activity, previous studies have quantified changes at several points during the recording, including at the onset of Ci-VSP activation and after repolarization ^26,27^. For WT GABA_A_Rs, however, repolarization from +50 mV to the –70 mV holding potential in our experiments induced tail currents resulting from increased membrane conductance during the membrane depolarization ^28^. This complicated the analysis of current amplitude changes after repolarization. We therefore quantified effects during the depolarization period from the onset of Ci-VSP activation, monitoring the effect of PI(4,5)P_2_ depletion on the outward agonist-evoked currents. To note, in GABA_A_R mutants and GlyRs that exhibited sensitivity to PI(4,5)P_2_ depletion, reductions in current amplitude were evident both during depolarization and after repolarization, but the quantification method was kept consistent throughout the study.

To eliminate contamination from endogenous activity in *Xenopus* oocyte recordings, Ci-VSP activation was first recorded using the same voltage command protocol in absence of agonist, thereby activating Ci-VSP without GABA_A_R or GlyR currents. Under these conditions, no inward current was detected, although a small outward current was sometimes observed in response to the membrane depolarization during Ci-VSP activation. These traces were subtracted from the corresponding agonist-evoked recordings to obtain isolated GABA_A_R or GlyR currents. Resulting traces were normalized to the peak of the outward current measured during the onset of depolarization, excluding the first 8 ms to avoid contamination from membrane capacitance peaks. Normalization was performed by dividing each current amplitude (sampled at 1-ms intervals) by the peak value, and the resulting normalized currents were plotted against time. Current reduction was quantified as the difference between the peak and steady-state current at the end of depolarization and expressed as a percentage.

### Analysis of agonist dose-response

Responses to GABA or glycine were normalized as *I_norm_ = (I/I_max_)*, where *I* represents the peak current amplitude at a given agonist concentration and I_max_ represents the maximal current evoked by the highest agonist concentration for each individual oocyte. Normalized responses were pooled and plotted as mean ± SEM. Dose-response data were fitted with the following equation:

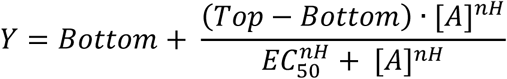

where Y is the peak current amplitude at agonist concentration [A], I_max_ is the maximal current, EC_50_ is the concentration required for half-maximal activation, and nH is the Hill coefficient. For all curve fit analyses, the Top parameter was constrained to 1.

### Analysis of patch clamp recording data

In outside-out patch clamp recordings with rapid perfusion, the 20–80% activation time was calculated as the time required for the current to rise from 20% to 80% of the peak amplitude. Deactivation and desensitization were characterized by time constants derived from double exponential function fittings to the decay phases following 1-ms and 1-s GABA applications, respectively. Weighted tau was calculated using the following equation:

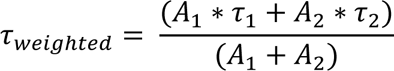

### Atomistic Molecular Dynamics (MD) simulations

Structural data were obtained from the Protein Data Bank (PDB), GABA_A_R (PDB ID:6I53) and GlyR (PDB ID:6UBS). Using the Modeller program ^29^, we modelled these structures with the mouse sequences corresponding to constructs used in electrophysiology experiments. For GABA_A_R, an α1K311N mutant structure was also generated. The channels used in the simulations were comprised of subunits which contained truncated intracellular domains between transmembrane helices M3 and M4 transmembrane helices (GABA_A_R α1, residues <u>328</u>–<u>381</u>, PEKPK<u>K</u>…<u>P</u>KKTFN; GABA_A_R β3, residues <u>312</u>–<u>415</u>, FGRGP<u>Q</u>…<u>P</u>DLTDV; GABA_A_R γ2L, residues <u>323</u>–<u>406</u>, HYFVS<u>N</u>…<u>R</u>IAKME; GlyR α1, residues <u>323</u>–<u>365</u>, RRKRR<u>H</u>…<u>P</u>PPAPS).

The MD simulation systems were set up using CHARMM-GUI web server ^30^. Each predicted channel complex was embedded into a lipid bilayer composed of 1,2-dipalmitoyl-sn-glycero-3-phosphocholine (DPPC) enriched with cholesterol at ∼30% (outer leaflet) and ∼25% (inner leaflet). The bilayer composition (inner leaflet) was adjusted to achieve a final enrichment of ∼10% PI(4,5)P_2_ for GABA_A_R systems and ∼8% for GlyR systems.

TIP3P model water ^31^ was used to solvate the bilayer complexes under a periodic boundary condition, and then an appropriate number of potassium and chloride ions replaced with water molecules to neutralize the system at an ionic strength of 150 mM.

We used the Particle Mesh Ewald method ^32^ to calculate electrostatic interactions. Chemical bonds involving hydrogen atoms other than water were treated as rigid using the LINCS algorithm ^33^. The SETTLE algorithm ^34^ was applied to the water model. The CHARMM force field ^35^ was employed for proteins, lipids, and ions. The time step was set to 2 fs.

The total energy of the prepared systems was minimized using the steepest descent method. Next, the system was equilibrated with a constant number of atoms, a fixed volume, and a temperature of 310 K. The system was further equilibrated with a constant number of atoms and a pressure of 1 bar at 310 K. For the production runs, the Parrinello-Rahman barostat ^36^ and the Nosé-Hoover thermostat ^37,38^ were used. We then saved snapshot structures at appropriate intervals from the trajectories for the analysis.

MD simulations were conducted by using GROMACS program ^39^. Trajectory visualization was performed using VMD ^40^, and structural figures were generated in PyMOL (The PyMOL Molecular Graphics System, Version 3.1.0 Schrödinger, LLC). Analysis of protein and lipids were performed using MDAnalysis ^41,42^ with Python scripts executed in Google Colaboratory.

### Analysis of PI(4,5)P_2_ contact probability in MD simulation

For all simulations, structural stability was assessed by calculating the root-mean-square deviation of the entire protein, using the initial frame as the reference. All simulations indicated the system equilibrated after the first 100 ns. For the following PI(4,5)P_2_ binding analysis, data were analyzed by omitting the initial 100 ns of each trajectory.

To characterize the interaction between residues of GABA_A_Rs or GlyRs and PI(4,5)P_2_ molecules, we performed a per-residue contact analysis using the MDAnalysis library ^41,42^. For GABA_A_Rs, residues R248, K311 (or N311), R312, and K390 were selected as reference points for distance calculations. For homomeric α1 GlyRs, because we lack the prior structural information regarding PI(4,5)P_2_ binding, we first identified putative binding residues from an initial screening of lipid binding events where we measured distances of PI(4,5)P_2_ phosphates to the α-carbon of residues that make up the entire protein (Extended Data Fig. 10b). For this initial screening in GlyRs, contact was defined as any distance ≤ 10 Å. For selected residues of GABA_A_Rs and identified residues of GlyRs, PI(4,5)P_2_ binding was further assessed by defining lipid contact as distances ≤ 5 Å between the phosphates of PI(4,5)P_2_ and the most terminal carbon (furthest carbon atom from the α-carbon) of residues. For each of the selected residues, binding events were quantified by classifying bound vs. unbound events and generating binarized time-series contact histories for all PI(4,5)P_2_ molecules across the sampled trajectory. From these time-series, we calculated three primary metrics to characterize PI(4,5)P_2_ binding: (1) contact probability, (2) max residence time, and (3) mean residence time. For (1) contact probability was defined as the fraction of the total simulation time (minus the initial 100 ns) during which a PI(4,5)P_2_ molecule was within the 5 Å distance of target residues. For (2) and (3), residence time was defined for each PI(4,5)P_2_ as the continuous contact events that occurred with each residue. Transitions between bound vs. unbound were identified by calculating the discrete derivative of the contact history, where a transition from 0 to 1 indicated a binding event and 1 to 0 indicating unbinding. The (2) max residence time is defined as the duration of the single longest continuous contact event that was recorded, representing the most stable binding. Finally, (3) mean residence time is defined as the average duration of continuous contact events.

Across three independent simulations, per phosphate analysis showed that the 1-phosphate (phosphodiester) contributed negligibly to the overall interaction of PI(4,5)P_2_ with the basic residues, whereas the 4-phosphate showed higher contact probability and the 5-phosphate the highest contact probability (Supplementary Fig. 2, and Supplementary Table 3).

### Statistical analysis

All data are presented as mean ± SEM unless otherwise indicated. Data analyses were performed using Igor Pro software (WaveMetrics Inc., USA). Statistical analyses were carried out in GraphPad software (Dotmatics, USA), with appropriate tests selected according to the experimental design.

## RESULTS

### Charge neutralization in the PI(4,5)P_2_-coordinating region confers sensitivity of GABA_A_Rs to acute PI(4,5)P_2_ depletion by Ci-VSP

We heterologously expressed mouse α1 (Fig. 1a), β3, and γ2L GABA_A_R subunits in *Xenopus* oocytes (hereafter referred to as GABA_A_R). To examine whether PI(4,5)P_2_ modulates wild-type (WT) GABA_A_R channel function, *Ciona intestinalis* voltage-sensing phosphatase (Ci-VSP) was co-expressed to induce depletion of endogenous membrane PI(4,5)P_2_ upon depolarization of the membrane potential (Fig. 1b) ^43^ and I_GABA_ were recorded upon depletion of PI(4,5)P_2_ in two-electrode voltage clamp (TEVC) mode. While inward I_GABA_ was evoked in response to 2.5 μM GABA at a holding potential of –70 mV, a 10-s depolarization step pulse was applied to activate the Ci-VSP. This depolarization reverses the direction of I_GABA_ from inward to outward. We therefore assessed the putative effect of depleting endogenous membrane PI(4,5)P_2_ on GABA_A_R channel activity by monitoring the outward I_GABA_ during the depolarization step (see Materials and Methods). The activity of WT GABA_A_Rs showed no difference over time, no matter whether Ci-VSP or the catalytically inactive Ci-VSP mutant (Ci-VSP^C363S^) was co-expressed and activated by depolarizing the membrane potential (Extended Data Fig. 1). These findings are consistent with a previous report demonstrating that PI(4,5)P_2_ depletion does not significantly alter I_GABA_ ^13^.

Extensive studies of PI(4,5)P_2_ regulation of ion channels have demonstrated that the spectrum of differential sensitivities to PI(4,5)P_2_ depletion reflect the different binding affinities of each channel ^8,44–46^. We therefore hypothesized that a strong affinity for PI(4,5)P_2_ in GABA_A_Rs prevents efficient release of PI(4,5)P_2_ from the channels during acute depletion of endogenous PI(4,5)P_2_ by Ci-VSP. In the cryo-electron microscopy (cryo-EM) structures, four positively charged amino acids (R248, K311, R312, and K390) of the α1 subunits form salt bridges with the negatively charged phosphates of the PI(4,5)P_2_ ^10^. We introduced charge-neutralizing mutations at these structurally identified binding sites (Fig. 1a) and assessed the mutational effects on the responses to PI(4,5)P_2_ depletions. α1K311N and α1R312L GABA_A_R mutants showed pronounced reductions in I_GABA_ following Ci-VSP activation (Fig. 1c–e and Extended Data Fig. 2b). In contrast, oocytes expressing α1R248Q GABA_A_Rs and α1K390Q GABA_A_Rs did not exhibit sensitivity to PI(4,5)P_2_ depletion (Fig. 1d,e and Extended Data Fig. 2a,c). α1R248Q GABA_A_Rs appear to require higher agonist concentrations (5 µM) to evoke currents comparable to WT (Fig. 1d,e and Extended Data Fig. 2a). Of note, α1K311N GABA_A_Rs displayed the highest sensitivity to PI(4,5)P_2_ depletion (Fig. 1d,e). α1K311N GABA_A_Rs showed a slight rightward shift of the GABA dose-response curves compared to WT GABA_A_Rs (Extended Data Fig. 5a,b) showing an approximately 1.4-fold in EC_50_ (WT, 10.73 ± 0.97 µM; α1K311N, 14.99 ± 1.99 µM) without alteration of the Hill coefficient (WT, 2.36 ± 0.13; α1K311N, 2.48 ± 0.16; Extended Data Fig. 5c). These results are comparable to previously reported properties of charge-neutralized mutants at the same position ^47^.

The effects of PI(4,5)P_2_ depletion by Ci-VSP were also examined using a paired pulse protocol (Extended Data Fig. 4a). Current responses to varying GABA concentrations (10 µM, 30 µM, and 1 mM GABA) were examined in oocytes co-expressing α1K311N GABA_A_R and Ci-VSP, first without VSP activation and then a second time subsequent to VSP activation (Extended Data Fig. 4b). The ratio of the second current amplitude (after Ci-VSP activation) to the first (without Ci-VSP activation) showed that depleting PI(4,5)P_2_ resulted in a significant reduction of current amplitude with 10 µM GABA (33.41 ± 6.014%; n = 4), but the effect became less apparent with higher GABA concentrations, 30 µM and 1 mM (8.77 ± 2.97% and 1.17 ± 0.044%, respectively; n = 5–15) (Extended Data Fig. 4c).

Together, these results suggest that neutralizing the positive charge of K311 weakened the tight binding of PI(4,5)P_2_.

### Caged lysine experiments indicate a critical role of PI(4,5)P_2_ binding in GABA_A_R channel activity

We infer that PI(4,5)P_2_ binds to WT GABA_A_Rs too tightly to be removed by the phosphatase activity of Ci-VSP and this high-affinity binding is mediated by K311. To prove this hypothesis, we used a recently developed approach that enables the optical control of PI(4,5)P_2_ binding to a single lysine residue to PI(4,5)P_2_ ^16^ (Fig. 2b). This method uses genetic code expansion technology ^48,49^ to site-specifically incorporate a caged lysine, hydroxycoumarin lysine (HCK), which can be converted to lysine upon UV light irradiation ^20^ (Fig. 2a). Previously, we successfully applied this approach in the mouse Kir2.1 potassium channel ^16^ and demonstrated PI(4,5)P_2_-dependent channel activity can be controlled by photon-induced uncaging of the unnatural amino acid ^50–55^.

We co-expressed the α1 subunit with incorporation of HCK at the position of K311 together with β3 and γ2L subunits (α1K311HCK GABA_A_R) to assess if UV-uncaging leads to the alteration of I_GABA_. Oocytes expressing α1K311HCK GABA_A_Rs were voltage-clamped at –70 mV and exposed to UV irradiation for 50–60 s during 2.5 µM GABA application. UV irradiation produced a 58.3 ± 3.4% increase in current amplitude, quantified as the change from the pre-irradiation baseline current to the peak current during irradiation (Fig. 2c). Serving as control, incorporation of HCK at lysine residue, K390, did not alter the current amplitude under the same conditions (Fig. 2d), consistent with our results showing no significant effect of PI(4,5)P_2_ depletion on the currents from α1K390Q GABA_A_Rs (Fig. 1d,e and Extended Data Fig. 2c). Oocytes lacking one or more components required for HCK incorporation produced only negligible GABA-evoked responses (Supplementary Fig. 1).

Next, we used the paired pulse protocol, to compare current amplitudes before and after UV irradiation across GABA concentrations of 10 µM, 30 µM, and 100 µM (Fig. 2e). Increases in current amplitude were observed at all three concentrations (10 µM, 1.85 ± 0.10-fold; 30 µM, 1.57 ± 0.08-fold; 100 µM, 1.17 ± 0.04-fold; n=5–15) (Fig. 2f). Notably, similar to our results from the α1K311N mutant co-expressed with Ci-VSP, uncaging effects were dose-dependent, with smaller changes at higher GABA concentrations (Fig. 2g).

Dose-response analysis of caged (α1K311HCK) mutants showed a slight rightward shift in the GABA dose-response curves compared to WT GABA_A_Rs (Extended Data Fig. 5a,b). Corresponding to this shift was an approximately 1.6-fold increase in EC_50_, respectively (WT, 10.73 ± 0.97 µM; α1K311HCK, 16.31 ± 0.40 µM), while the Hill coefficient was not significantly altered (WT, 2.36 ± 0.13; α1K311HCK 2.80 ± 0.12; Extended Data Fig. 5c).

To verify that the current increase observed upon uncaging of α1K311HCK GABA_A_Rs is mediated by the alteration of PI(4,5)P_2_ binding to the α1 subunit, we compared the sensitivity to PI(4,5)P_2_ depletion induced by Ci-VSP before and after uncaging (Fig. 3a,b). Prior to uncaging, α1K311HCK GABA_A_Rs displayed sensitivity to PI(4,5)P_2_ depletion, similar to the phenotype observed in α1K311N GABA_A_Rs and other charge-neutralized or charge-reversed mutants (Fig. 1c and Extended Data Fig. 3). Following uncaging, current reduction during Ci-VSP activation was no longer observed, indicating that uncaging restored the channel activity to a WT-like phenotype (Fig. 3b–d). These results suggest that the increase in current magnitude upon UV irradiation was due to the tighter binding of PI(4,5)P_2_ to the channel.

### PI(4,5)P_2_ binding is required for rapid activation of GABA_A_Rs at physiological ligand concentration

Our TEVC results showed that the reduction in current following PI(4,5)P_2_ depletion and the increase in current following K311 uncaging were more pronounced at lower than at higher agonist concentrations, which suggests that PI(4,5)P_2_ specifically affects GABA_A_R channel kinetics only at submaximal conditions. However, activation kinetics of GABA_A_Rs at higher agonist concentrations have been reported to occur on the millisecond timescale and time resolution in TEVC mode is not optimal to catch such rapid kinetics ^56,57^. Therefore, we performed outside-out patch clamp recordings using piezo-controlled fast agonist application system ^58–60^.

In outside-out patch clamp mode, WT GABA_A_R currents were recorded in response to the physiological GABA concentration of 1 mM ^61–63^, at a holding potential of −70 mV under symmetrical chloride conditions, resulting in inward I_GABA_. The 20–80% activation time for WT, measured from responses to 1-ms applications of 1 mM GABA, was 0.43 ± 0.02 ms (n = 8; Fig. 4a,c,d). Current decay from the same protocol was fitted with a double-exponential function, yielding a fast time constant (τ_fast_) of 7.01 ± 1.14 ms, slow time constant (τ_slow_) of 164.60 ± 23.47 ms and a weighted time constant (τ_weighted_) of 48.65 ± 9.60 ms (n = 8; Fig. 4a and Extended Data Fig. 6e). Desensitization kinetics for WT were analyzed from responses to 1-s GABA application protocol and fitted with a double exponential function, yielding τ_fast_ = 9.41 ± 1.47 ms, τ_slow_ = 387.40 ± 50.73 ms, and τ_weighted_ = 123.30 ± 25.02 (n = 8; Extended Data Fig. 6a,d). These parameters of kinetics in WT channels are comparable to those reported in previous studies, which recorded GABA_A_Rs with the same subunit composition under slightly different experimental conditions ^64^.

**Figure 4.**
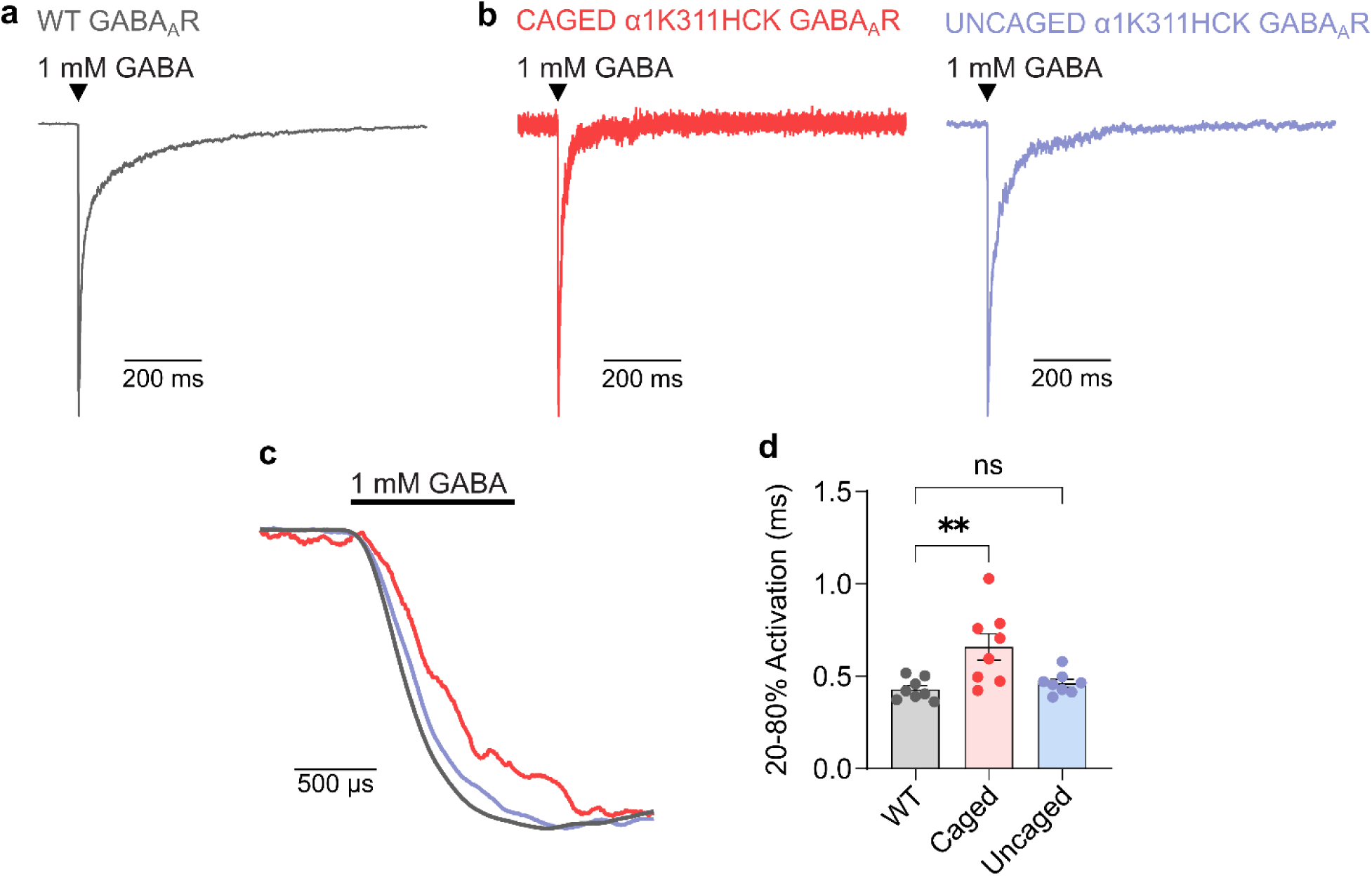
PI(4,5)P_2_ binding is required for rapid activation of GABA_A_Rs at physiological ligand concentration. Normalized representative outside-out patch-clamp recording traces of **(a)** WT and **(b)** α1K311HCK GABA_A_Rs in caged state (light red) and uncaged state (violet blue). **(c)** Magnified superimpose of normalized outside-out patch-clamp recording traces of WT (gray), α1K311HCK GABA_A_Rs in caged state (light red) and α1K311HCK GABA_A_Rs in uncaged state (violet blue) in (a) and (b) from 101 ms to 103.4 ms time scale. **(d)** Comparison of 20–80% activation time between α1K311HCK GABA_A_Rs caged and uncaged compared to WT GABA_A_Rs (20–80% activation time WT = 0.43 ± 0.02 ms; caged = 0.66 ± 0.07 ms; uncaged = 0.46 ± 0.02 ms; n = 8). Statistical significance was assessed with Kruskal-Wallis test followed by Dunn’s post hoc test (p = 0.008 for WT vs. Caged; p = 0.832 for WT vs. Uncaged; n = 8). Data represents mean ± SEM.

Next, we performed outside-out patch clamp recordings in α1K311HCK GABA_A_Rs. Responses to a 1-ms GABA application followed by a 1-s GABA application protocol were analyzed in the caged state and uncaged state in the same patch (Fig. 4b,c and Extended Data Fig. 6b,c). Two series of recordings were intervened by 2-min irradiation with UV light. Strikingly, the 20–80% activation rise time was accelerated by approximately 1.4-fold upon uncaging (caged = 0.66 ± 0.07 ms; uncaged = 0.46 ± 0.02 ms; n=8; Fig. 4b–d). The 20–80% activation time after uncaging was comparable to that of WT (Fig. 4d). Apart from 20–80% activation time, there were no differences in deactivation or desensitization kinetics, nor in residual currents between WT GABA_A_Rs, caged α1K311HCK GABA_A_Rs, and uncaged α1K311HCK GABA_A_Rs (Extended Data Fig. 6d–e), although we observed variability in the parameters.

These results show that the activation rise time of α1K311HCK GABA_A_Rs was accelerated upon uncaging, strongly suggesting that PI(4,5)P_2_ binding mediated by K311 is required for rapid GABA_A_Rs activation.

### PI(4,5)P_2_ binds stably to K311 and R312 of GABA_A_Rs

The results from our electrophysiology experiments suggest that the lysine residue K311 plays a critical role in mediating the high binding affinity for PI(4,5)P_2_ in GABA_A_Rs. While structural studies have defined how PI(4,5)P_2_ binds to GABA_A_Rs, the steric interactions underlying the high-binding affinity remained unclear. To investigate the dynamics of PI(4,5)P_2_ interaction with K311 and the other identified residues, we performed atomistic molecular dynamics (MD) simulations. For this, we used α1β3γ2L GABA_A_R modelled with mouse sequences corresponding to subunit constructs used in the electrophysiology experiments (Fig. 5a).

**Figure 5.**
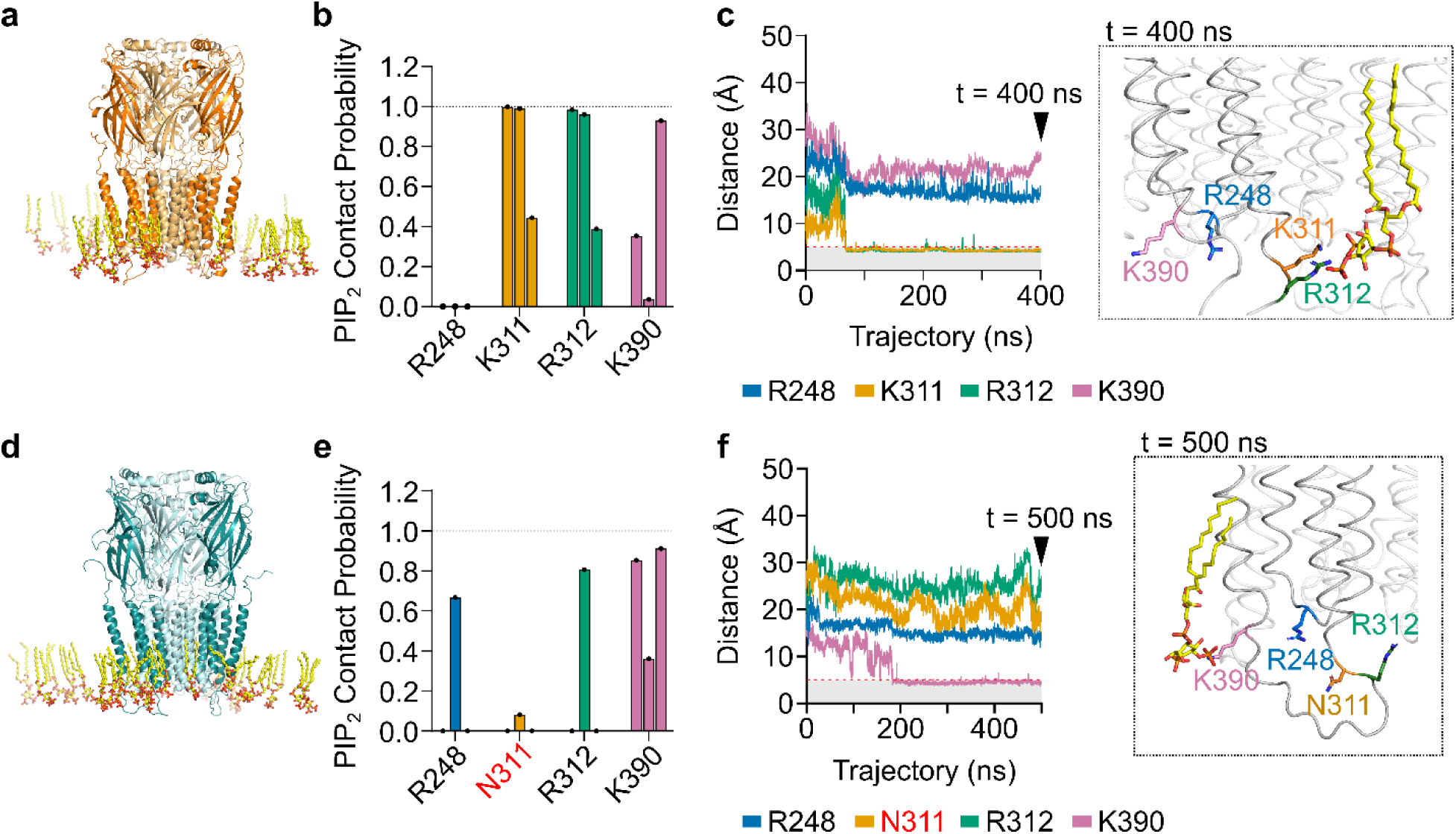
Neutralizing the positive charge at K311 weakens PI(4,5)P_2_ binding but does not abolish the interaction. **(a)** Representative structure of WT GABA_A_Rs in the presence of 10% PI(4,5)P_2_. **(b)** PI(4,5)P_2_ contact probability was analyzed as the fraction of the trajectory during which any PI(4,5)P_2_ molecules were in < 5 Å proximity to the most terminal carbon atom residues (furthest from Cα). Data were obtained from three independent simulations (400–500 ns); each bar represents a single repeat. **(c)** P-C distances (from 5-phosphate) to each residue are shown. A snapshot at was taken at the end of the trajectory (t = 400 ns). **(d)** Representative structure of α1K311N GABA_A_Rs + 10% PI(4,5)P_2_. **(e)** PI(4,5)P_2_ contact probability and **(f)** P-C distances (from the 5-phosphate to terminal carbon of residues) are shown similar to (b-c). A snapshot was taken at t = 500 ns to show the final structure of the PI(4,5)P_2_ binding in α1K311N. Data were obtained from three independent simulations (400–500 ns); each bar represents a single repeat in **(e)**.

In WT GABA_A_R, PI(4,5)P_2_ binding behavior was largely consistent with cryo-EM structures, primarily coordinated by residues of the α1 subunit ^10,11^. Of the four basic residues, K311 and R312 accounted for most of the contacts across all simulations (Fig. 5b). Following the interaction of one PI(4,5)P_2_ molecule, K311 and R312 exhibited strong interaction with the PI(4,5)P_2_, demonstrating long residence times within minimum distances of ≤ 5 Å (Fig. 5c). Indeed, residence time calculations revealed that while many binding events consisted of short-lived < 5 ns binding events, K311 and R312 supported most sustained binding (∼44.83 ns and ∼113.08 ns, respectively). In contrast, K390 showed slightly briefer contacts (∼25.88 ns) and R248 showed no contact events at all (Supplementary Table 3). Of note, although K390 still exhibited rather high PI(4,5)P_2_ contact probability (Fig. 5b), the PI(4,5)P_2_ molecules that bind K390 were distinct from those interacting with K311 and R312 (Extended Data Fig. 7). Despite the observed variability amongst the replicated simulations, these results indicate that of the four residues implicated as PI(4,5)P_2_ binding sites, K311 and R312 show the most stable binding. This result is consistent with our electrophysiological data demonstrating that charge-neutralized mutations of K311 and R312, in contrast to R248 and K390, confers the GABA_A_Rs sensitive to acute PI(4,5)P_2_ depletion.

### Neutralizing K311 reduces the stability of PI(4,5)P_2_ binding

We then performed atomistic MD simulations on the α1K311N GABA_A_Rs (Fig. 5d). While the mutation of K311 markedly reduced its direct interactions with PI(4,5)P_2_, the lipid remained interacting with other residues, R248, R312, and K390 (Fig. 5e,f and Supplementary Table 4). Analysis of the simulation repeats revealed a shift in the binding profile; notably, residue R248 maintained a robust interaction, in one repeat even with a contact probability of ∼0.83 and a maximum residence time of ∼218.15 ns. In contrast, R312 exhibited reduced binding stability compared to the WT channel; while the contact probability remained relatively consistent at ∼0.61 (WT, ∼0.72), the residence time decreased to ∼75.64 ns from ∼113.08 ns of WT channels. Interaction at K390 remained largely comparable to WT, showing a contact probability of ∼0.40 and a residence time of ∼35.86 ns (WT, ∼0.51 and ∼25.88 ns, respectively). Collectively, a neutralizing mutation of K311 does not fully abolish the interaction with PI(4,5)P_2_ in GABA_A_Rs, but results in a qualitatively less stable interaction profile characterized by reduced residence times at key residues.

### PI(4,5)P_2_ modulates GlyR channel activity

Glycine receptors (GlyRs) share both structural and functional similarities with GABA_A_Rs. In GlyR α1 subunits, several positively charged residues are conserved at positions corresponding to the PI(4,5)P_2_ binding sites previously identified in cryo-EM structures of GABA_A_Rs, most notably the K311 of GABA_A_R α1 subunit ^10,11^. Of reported GlyR structures (Supplementary Table 5), the structure of a zebrafish homopentameric α1 GlyRs reveals several lipid densities with one identified within the inner leaflet at the M3-M4 interface ^65^. However, these lipid densities were only modelled as partial phospholipids, and whether PI(4,5)P_2_ binds to or modulates GlyR channel activities remained unclear. We expressed α1 GlyRs with Ci-VSP (Fig. 6a) in *Xenopus* oocytes and currents in response to 20 µM glycine (I_glycine_) were recorded. I_glycine_ during a depolarizing pulse (+50 mV, 10-s) showed clear decline upon PI(4,5)P_2_ depletion whereas co-expression of the catalytically inactive Ci-VSP^C363S^ mutant did not yield such decline of I_glycine_ (Fig. 6a–c). In addition, we assessed the effect of depleting PI(4,5)P_2_ prior to agonist application in the paired pulse protocol. Experiments at varying glycine concentrations (50 µM, 100 µM, and 1 mM) showed a dose-dependent effect, with progressively smaller reductions at higher agonist concentrations (50 µM, 46.05 ± 4.34%; 100 µM, 16.82 ± 3.71%; and 1 mM, 14.78 ± 3.031%; n = 7-14) (Extended Data Fig. 8).

**Figure 6.**
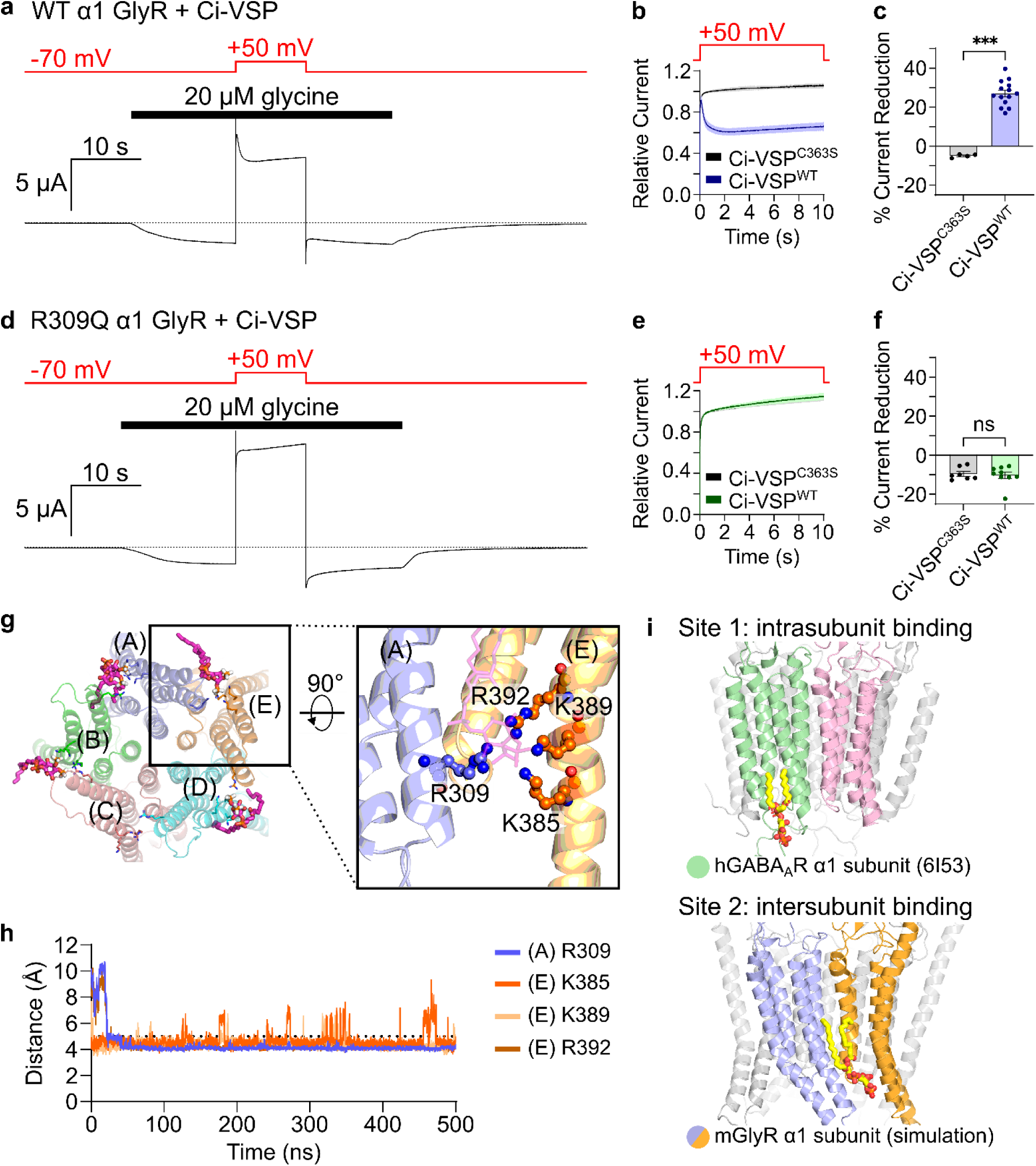
Homopentameric α1 GlyR exhibits sensitivity to acute PI(4,5)P_2_ depletion by Ci-VSP. **(a)** Representative TEVC traces of WT GlyR co-expressed with Ci-VSP. **(b)** Summarized time course plot of normalized outward I_glycine_ during Ci-VSP activation. **(c)** Percent current reduction compared between GlyRs co-expressed with Ci-VSP^WT^ and Ci-VSP^C363S^. (d) Representative TEVC trace of α1R309 GlyR co-expressed with Ci-VSP. **(e-f)** Summarized time course plot of normalized outward I_glycine_ **(e)** and percent current reduction compared between α1R309 GlyRs co-expressed with Ci-VSP^WT^ and Ci-VSP^C363S^ **(f)**. **(g)** Bottom-view snapshot from atomistic MD simulation demonstrating multiple PI(4,5)P_2_ molecules bound to the GlyR α1 subunit. Expanded side view highlights the residues, with distance plots between the P5-phophsate atom and the identified residues shown below. **(i)** Comparison of PI(4,5)P2 binding sites in cryo-EM structure of human GABAAR (PDB ID: 6I53) and GlyRs from simulation. Plots of current decay during Ci-VSP activation (b and e) shows means ± SEM (shading). Statistical significance of percent current reduction (c and f) was assessed using Mann-Whitney U test (WT, p = <0.0001; α1R309Q, p = 0.837; n = 7–14 oocytes from 2–3 batches). Data represents mean ± SEM.

We next tested the effect of mutating the residue R309, which corresponds to the K311 of the GABA_A_R α1 subunit. A R309Q mutation abolished the sensitivity to acute PI(4,5)P_2_ depletion (Fig. 6d–f). Dose-response analysis of α1R309Q GlyRs showed comparable channel properties as WT α1 GlyR (Extended Data Fig. 9).

We finally performed atomistic MD simulations, using the zebrafish homopentameric α1 GlyR structure (PDB ID: 6UBS) as a template to generate a mouse homopentameric α1 GlyR. Unlike the previous study of cryo-EM structure analysis showing a lipid density located in the intrasubunit region of the α subunit ^65^, binding of PI(4,5)P_2_ was observed mostly in the intersubunit regions (Fig. 6g–i and Extended Data Fig. 10). Indeed, binding of PI(4,5)P_2_ to GlyR was coordinated by the R309, which corresponds to K311 in the α1 subunit of GABA_A_Rs, but also by residues of the M4 helix of the neighboring subunit including K385, K389, and R392 (Fig. 6g,h; Extended Data Fig. 10). Thus, the intersubunit binding profile of PI(4,5)P_2_ in GlyRs contrasts with the intrasubunit coordination of GABA_A_Rs (Fig. 1a and Fig. 6i) ^10,11^.

## DISCUSSION

Our study unravels a previously unappreciated functional role of PI(4,5)P_2_ in GABA_A_R channel activity. Combining state-of-the-art electrophysiology and MD simulations with a recently developed caged lysine technology that allows alteration of PI(4,5)P_2_ binding affinity in the same protein, we demonstrate that the high-affinity binding of PI(4,5)P_2_ facilitates channel activity of GABA_A_Rs for the following pieces of evidence: (1) targeted mutations of specific basic amino acid residues, particularly K311, previously identified as a PI(4,5)P_2_ binding site ^10,11^, rendered GABA_A_Rs sensitive to PI(4,5)P_2_ depletion, which led to compromised channel function. (2) photocaging K311, which allowed the optical control of PI(4,5)P_2_ binding, revealed that uncaging results in not only an augmentation of channel activity but also accelerated the activation of GABA_A_Rs. (3) neutralizing the high-affinity PI(4,5)P_2_ binding site K311 did not abolish overall PI(4,5)P_2_ interaction, but only weakened its binding affinity, which strongly suggests that fast activation of GABA_A_Rs requires rigid docking of PI(4,5)P_2_. Thus, we propose a new paradigm for PI(4,5)P_2_ as a constitutive co-factor for proper GABA_A_R function in fast synaptic neurotransmission. Finally, as GlyRs also exhibit PI(4,5)P_2_-dependent channel activity, the requirement of PI(4,5)P_2_ may constitute a fundamental feature across inhibitory pLGICs.

The very high affinity of PI(4,5)P_2_ binding to GABA_A_Rs may well explain why this regulatory requirement has remained elusive in previous studies ^10,13^. Consistently, our data reveal that in WT channels, PI(4,5)P_2_ is bound so tightly that the channel activity remains unaffected throughout acute depletions of surrounding PI(4,5)P_2_. Only by reducing the binding affinity through charge-neutralizing mutagenesis, we were able to reveal effects of PI(4,5)P_2_ on GABA_A_R channel activity. Furthermore, the use of photocaging technology allowed us to demonstrate within the same population of receptors that GABA_A_R currents depend on interaction of the receptors with PI(4,5)P_2_.

Structural studies of the GABA_A_Rs resolving its open state conformation used to be hampered by the rapid gating kinetics of the receptor, which occur far faster than the typical window for cryo-EM sample preparation ^66–68^. Recently, this limitation was overcome by optimizing cryo-EM techniques to facilitate the capture of structures at millisecond timescales ^69^. Surprisingly, the resolved open state structure was devoid of PI(4,5)P_2_. This finding led the authors to claim that PI(4,5)P_2_ binding would rather inhibit channel activity. However, the study used a chimeric construct with the intracellular M3-M4 loop adjacent to K311, replaced by a shorter linker based on a bacterial homolog (SQPARAA). Notably, previous reports of chimeric GABA_A_Rs that also modified the M3-M4 loop with the same bacterial linker exhibited a gain-of-function phenotype^47^. Hence, the resolved open state structure of the chimeric constructs lacking PI(4,5)P_2_ molecules may not reflect a relief from an inhibition by PI(4,5)P_2_, but may rather exhibit an unexpected gain-of-function outcome. We assume that PI(4,5)P_2_ binds to GABA_A_Rs in a state-independent manner because both the enzyme-mediated depletion and the light-induced binding of PI(4,5)P_2_ affected GABA_A_R activity, whether ligand was bound (Fig. 1 and Fig. 3) or not (Fig. 2, Fig.4, and Extended Data Fig. 4). We hypothesize that M3-M4 loop containing the PI(4,5)P_2_ binding region is critical for channel activation and disrupting this interaction or remodeling this region leads to a loss-of-function or gain-of-function in GABA_A_Rs.

In heteromeric GABA_A_Rs, PI(4,5)P_2_ binds within an intrasubunit binding profile, whereas our simulations of homomeric GlyRs revealed an intersubunit binding profile, where PI(4,5)P_2_ sits between the M4 helix of one subunit and the M3 helix of the adjacent subunit. This intersubunit lipid binding profile has been previously observed in other members of the pLGIC family, suggesting it may represent the canonical binding profile of lipids ^6,70–72^. Recently, the honeybee GABA RDL receptor structure, a homopentameric channel, also revealed intersubunit PI(4,5)P_2_ binding ^73^. Collectively, these structural observations suggest that differences in how PI(4,5)P_2_ binds to pLGICs may depend homomeric versus heteromeric assembly. It will be intriguing to test whether the diverse binding profiles of PI(4,5)P_2_ may yield functional significance.

Our functional results strongly suggest K311 is critical for the PI(4,5)P_2_-mediated facilitation of GABA_A_Rs activation. These findings are consistent with previous studies where charge-switch and charge-neutralizing mutations were made at the K312 of the human α1 subunit (corresponding to the present K311 of mouse GABA_A_R) ^47^. Using partial agonists, the K312E mutant primarily affected gating efficacy rather than agonist binding affinity. Given that this residue also lies on the intracellular face of the transmembrane domain, distant from the agonist-binding site located in the extracellular domain, K311 may be fundamental for the pore opening mediated by the stabilization of the transmembrane helices through PI(4,5)P_2_ binding.

Both neutralizing K311 and caging K311 reduced GABA_A_R current upon PI(4,5)P_2_ depletion, whereas uncaging K311 increased it significantly. Ultra-rapid perfusion experiments with outside-out patch clamp experiments resolved that caging of K311 significantly delayed the transition of receptors from its closed to its active state, whereas uncaging K311 accelerated the activation rise time to WT-like speed. This suggests that PI(4,5)P_2_ acts as a facilitator that lowers the energetic barrier for pore opening ^74^. We hypothesize that PI(4,5)P_2_ stabilizes a preactive conformation that is important to enable the precision of fast synaptic transmissions. Of note, although we observed no significant differences in desensitization kinetics, due to some technical limitations of the current study (e.g. cell damage from UV exposure; see Materials and Methods), we may not completely exclude that PI(4,5)P_2_ is also involved in the desensitization gating of the channels. More detailed analysis such as single channel recordings will be necessary to determine which state transition PI(4,5)P_2_ regulates in the gating scheme of GABA_A_Rs.

Our data demonstrate that K311 is essential for the binding of PI(4,5)P_2_, which in turn facilitates the rapid activation that is the characteristic hallmark of synaptic GABA_A_Rs. Given that fast gating is a physiological requirement for synaptic transmission, it is noteworthy that structural evidence of PI(4,5)P_2_ has only been observed in synaptic isoforms (α1/2/3-containing GABA_A_Rs) ^10–12,75^. In striking contrast, extrasynaptic GABA_A_Rs, often composed of α4 or α6 subunits, contain an asparagine at the corresponding K311 site and in a recently reported structure of an α4-containing GABA_A_R, no PI(4,5)P_2_ was found bound ^76^. Therefore, it is plausible to hypothesize that unlike synaptic GABA_A_Rs where rapid gating is physiologically essential, extrasynaptic GABA_A_Rs may not necessitate such regulation.

The integration of structural and functional data with human genome databases may hint at a prediction for pathophysiological consequences of disrupted PI(4,5)P_2_ binding. Indeed, a variant in the human *GABRA1* gene (p.Lys339Glu), which corresponds to the mouse K311 residue, was listed in the ClinVar database (Accession: NC_000005.10:g.161895824A>G) with links to developmental and epileptic encephalopathy 19 (DEE19). Predictive algorithms (PolyPhen score = 0.998, SIFT = 0.00) indicate a likely deleterious effect by this mutation. Although not directly confirmed with patients suffering from DEE19, our data suggests that disrupting the electrostatic interaction of PI(4,5)P_2_ at this lysine residue site will result in delayed activation due to reduced PI(4,5)P_2_ affinity. With the growing recognition of PI(4,5)P_2_ as critical regulators of numerous ion channels, it has become clear that disrupted channel-PI(4,5)P_2_ interactions, either by channel dysfunction or phosphoinositide imbalance, represent a fundamental pathogenic mechanism in various channelopathies ^77^. For GABA_A_Rs, a disruption of the PI(4,5)P_2_ interaction may manifest as an imbalance between excitatory and inhibitory activity, eventually leading to conditions such as epilepsy.

In summary, by integrating structural, computational, and electrophysiological analyses, we establish that PI(4,5)P_2_ has functional roles in inhibitory pLGICs. Our findings uncover the mechanistic basis for the apparent insensitivity of WT GABA_A_Rs to PI(4,5)P_2_ depletion and show that PI(4,5)P_2_ is essential for supporting the channel activity of both GABA_A_Rs and GlyRs. Taken together, we propose PI(4,5)P_2_ as a critical endogenous co-factor that enables the proper channel function of inhibitory pLGICs. Moreover, in this study we utilized the unique property of caged lysine technology that allows the manipulation of PI(4,5)P_2_ binding affinity in the same protein, and revealed a previously inaccessible dynamics of PI(4,5)P_2_ binding in GABA_A_Rs. Our findings provide a foundation for future efforts to target PI(4,5)P_2_-GABA_A_R interactions as a means to fine-tune inhibitory signaling in health and disease.

## Acknowledgements

We thank S. Sugimoto for his help during the initial phase of this study and Dr. Byung-Chang Suh for his insightful advice during the study. We also thank Dr. Paul Brehm for his critical reading of the manuscript. This work was supported by the Center for Medical Research and Education at the Graduate School of Medicine, The University of Osaka and by World Premier International Research Center Initiative (WPI), Ministry of Education, Culture, Sports, Science and Technology (MEXT), Japan. Funding was provided by the Japan Society for the Promotion of Science (JSPS), Grant-in-Aid for JSPS Fellows (JP24KJ1563 to R.M., JP22KF0249 and JP22F22384 to Y.O. and R.T.A.), Grant-in-Aid for JSPS KAKENHI (JP22H02804 and JP25K02442 to Y.O., Y.Y., and T.K.), Grant-in-Aid for JSPS Exploratory Research (JP21K19350 to Y.O.), Japan Science and Technology (JST) FOREST Program (JPMJFR225Z to T.K.), and MEXT, Grant-in-Aid for Transformative Research Areas A (JP25H01334 to Y.O.), Grant-in-Aid for Transformative Research Areas B (JP20H05791 to Y.O.), and Grant-in-Aid for Scientific Research on Innovative Areas (JP15H05901 to Y.O.). Additionally, financial support was received from Mitsubishi Foundation and the Nakatani Foundation.

## Author informsation

These authors contributed equally: Risa Mori-Kreiner, Rizki Tsari Andriani

## Contributions

T.K., N.K., and Y.O. conceptualized and supervised the entire project. R.M., R.T.A., J.Z., and T.S. designed and performed electrophysiology experiments. N.M. and Y.Y. designed and performed computational experiments. R.M., R.T.A., and T.S. analyzed the data. R.M., R.T.A., T.S., T.K., N.K., and Y.O. contributed to the data interpretation and manuscript preparation. R.M., R.T.A., T.K., and Y.O. wrote the manuscript with input from all authors. All authors edited and approved the paper.

## Ethics declarations

### Disclosure for AI usage

During the preparation of this work, the authors acknowledge the usage of Google Gemini 1.5 and 3 Flash to assist with code editing and debugging in Python scripts for data analysis. After usage, the authors reviewed, validated and edited the code, taking full responsibility for the integrity of the generated output.

## Data availability

Extended data

Supplementary information

Source data

---

**Extended Data Figure 1.**
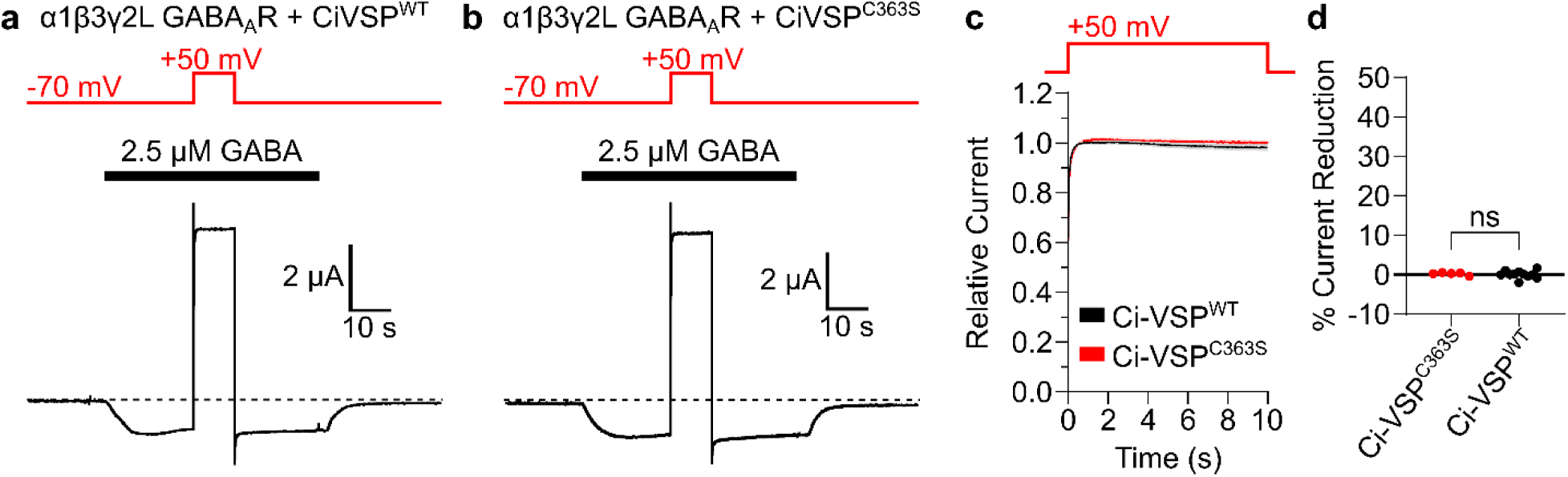
WT GABA_A_Rs are resistant to acute PI(4,5)P_2_ depletion by Ci-VSP. **(a)** Representative current trace of WT GABA_A_R co-expressed with Ci-VSP (top) and Ci-VSP^C363S^ mutant (bottom). **(b)** Time course plot comparing normalized outward I_GABA_ of GABA_A_R variants during the 10 s depolarization step pulse. Plot shows the group means ± SEM (shading); n = 5–11 oocytes per group from 2–3 batches. **(c)** Comparison of percent (%) current reductions between WT GABA_A_R co-expressed with Ci-VSP vs Ci-VSP^C363S^ mutant. Normal distribution was not assumed for the datasets and statistical significance was assessed by Mann-Whitney U test (p = 0.510). Data represents mean ± SEM.

**Extended Data Figure 2.**
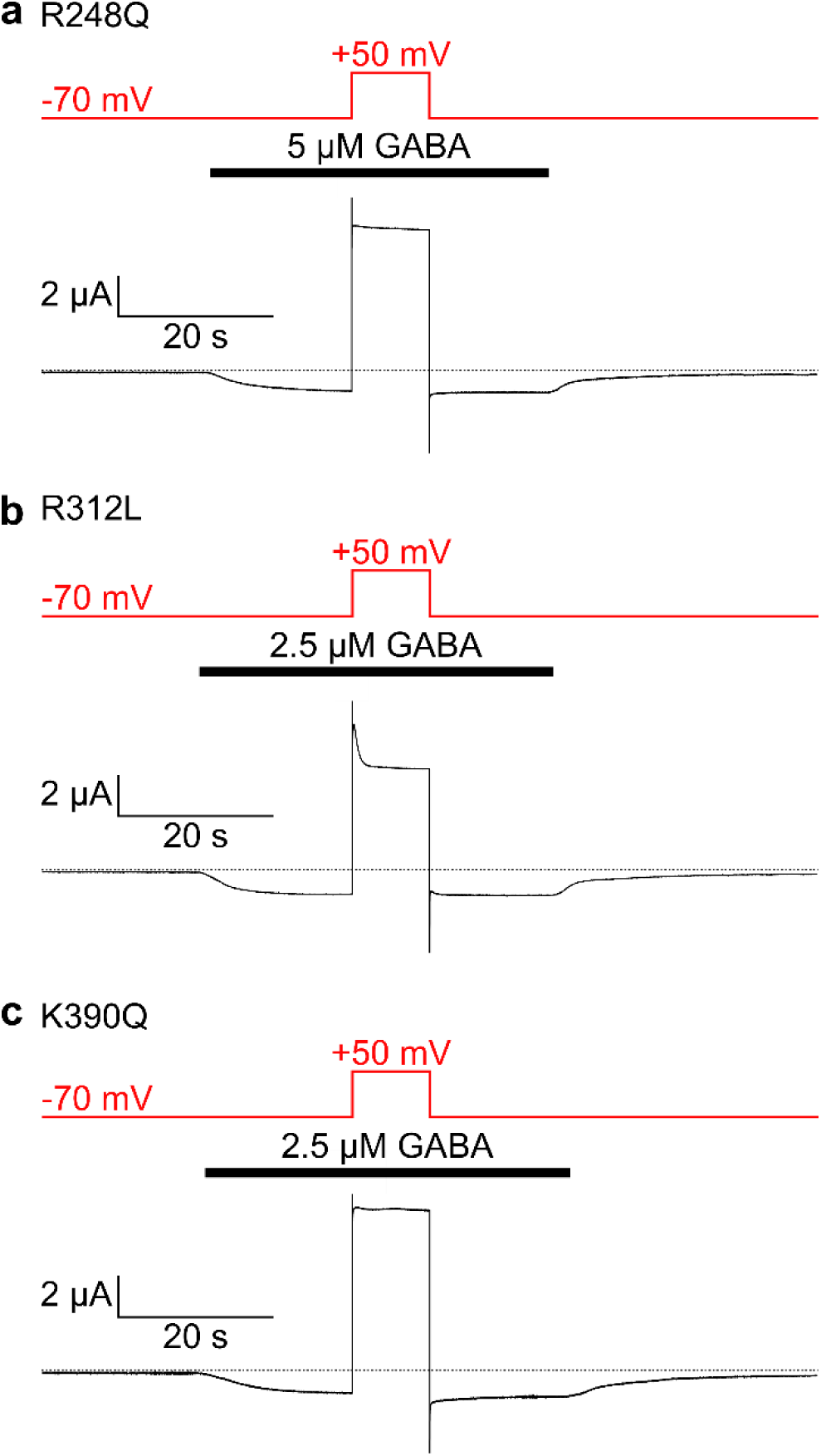
Neutralizing positive charges of identified PI(4,5)P_2_ binding sites result in varying degrees of sensitivity to acute PI(4,5)P_2_ depletion by Ci-VSP. **(a)** Representative current trace of α1R248Q GABA_A_R co-expressed with Ci-VSP. **(b)** Representative current trace of α1R312L GABA_A_R co-expressed with Ci-VSP. **(c)** Representative current trace of α1K390Q GABA_A_R co-expressed with Ci-VSP.

**Extended Data Figure 3.**
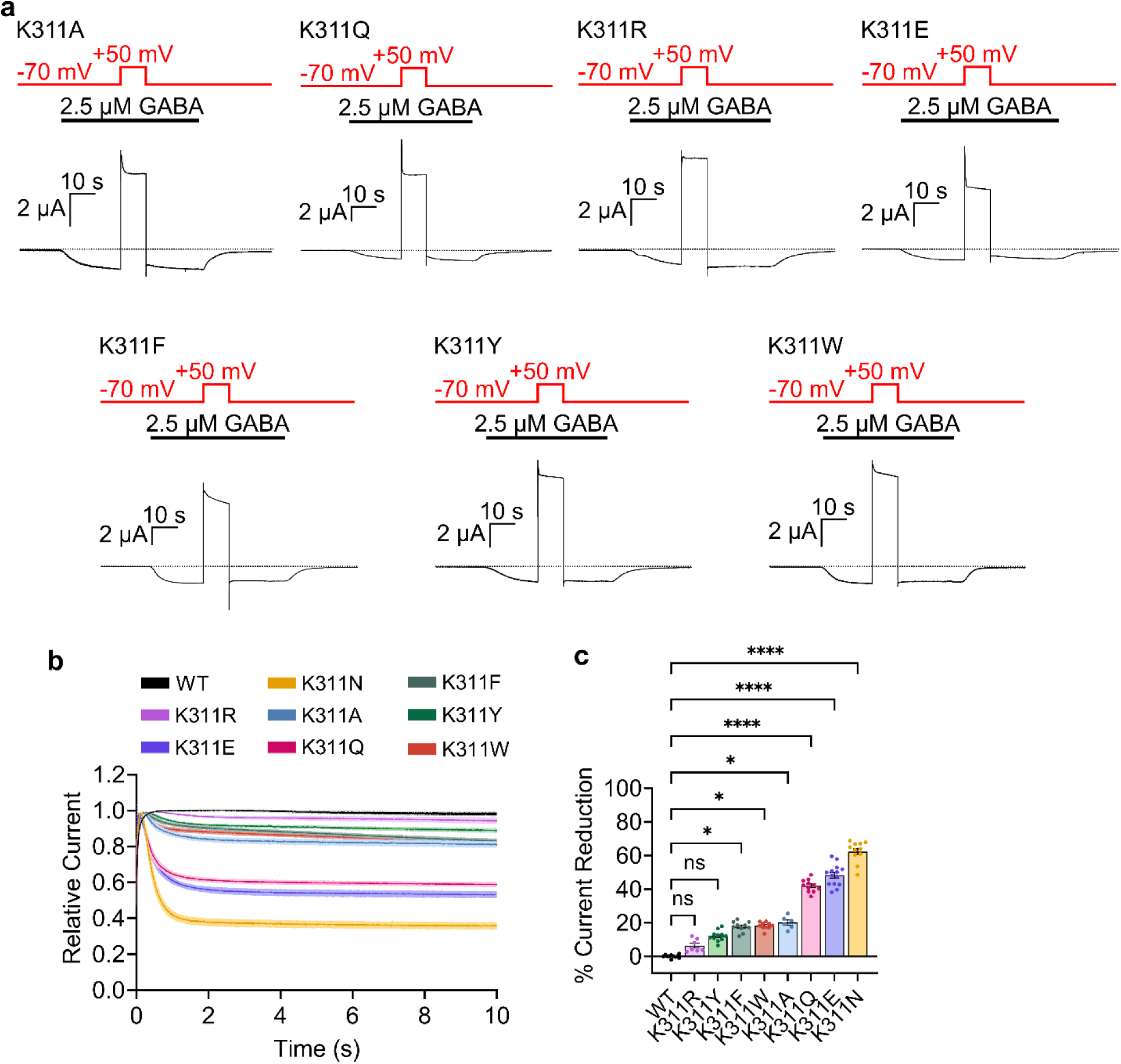
Substitution of residue K311 with different amino acids varying in charge, polarity, and size results in varying degrees of sensitivity to acute PI(4,5)P_2_ depletion by Ci-VSP. **(a)** Representative current traces of K311(α1) mutants mutated to alanine (A), arginine (R), glutamine (Q), glutamate (E), phenylalanine (F), tryptophan (W), and tyrosine (Y) co-expressed with Ci-VSP. **(b)** Time course plot comparing normalized outward I_GABA_ of GABA_A_R variants during the 10 s depolarization step pulse. Plot shows the group means ± SEM (shading). **(c)** Comparison of percent (%) current reductions across GABA_A_R variants during Ci-VSP activation (compared with WT and α1K311N mutant data from Figure 1 and Extended Data Figure 1). Statistical significance was assessed using Kruskal-Wallis test followed by the Dunn’s post hoc test (WT vs. α1K311A, p = 0.022; WT vs. α1K311Q, p = <0.0001; WT vs. α1K311R, p = >0.999; WT vs. α1K311E, p = <0.0001; WT vs. α1K311F, p = 0.019; WT vs. α1K311W, p = 0.012; WT vs. α1K311Y, p = 0.365). Data represents mean ± SEM.

**Extended Data Figure 4.**
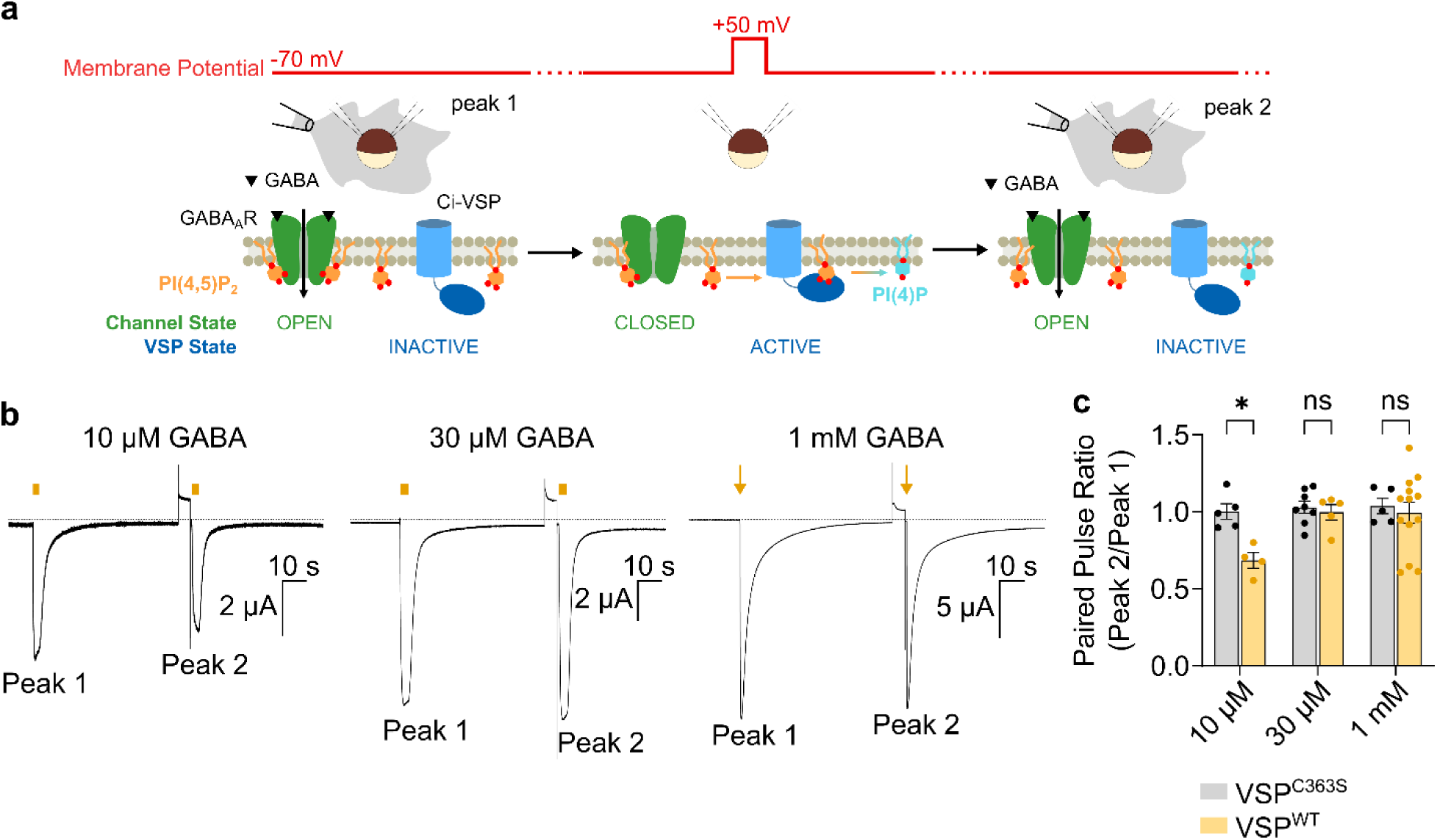
VSP activation in the unliganded state of α1K311N GABA_A_Rs result in reduced channel opening. **(a)** Schematic overview of the paired pulse TEVC experiment. Membrane potential settings indicated in red; GABA_A_R channel state (closed/open) indicated in green; VSP state (inactive/active) indicated in blue. **(b)** Representative traces of α1K311N GABA_A_R co-expressed with Ci-VSP; I_GABA_ with/without Ci-VSP activation at varying GABA concentrations (top, 10 μM GABA; middle, 30 μM GABA; bottom, 1 mM GABA). GABA applications are indicated by bars or arrows. **(c)** Summary of paired pulse ratios compared between α1K311N GABA_A_R co-expressed with Ci-VSP and Ci-VSP^C363S^ mutant at different GABA concentrations. Statistical significance was assessed using ordinary two-way ANOVA followed by Bonferroni’s post hoc test (10 µM, p = 0.029; 30 µM, p = >0.999; 100 µM, p = >0.999; n = 4–13 oocytes from 2–3 batches). Data represents mean ± SEM.

**Extended Data Figure 5.**
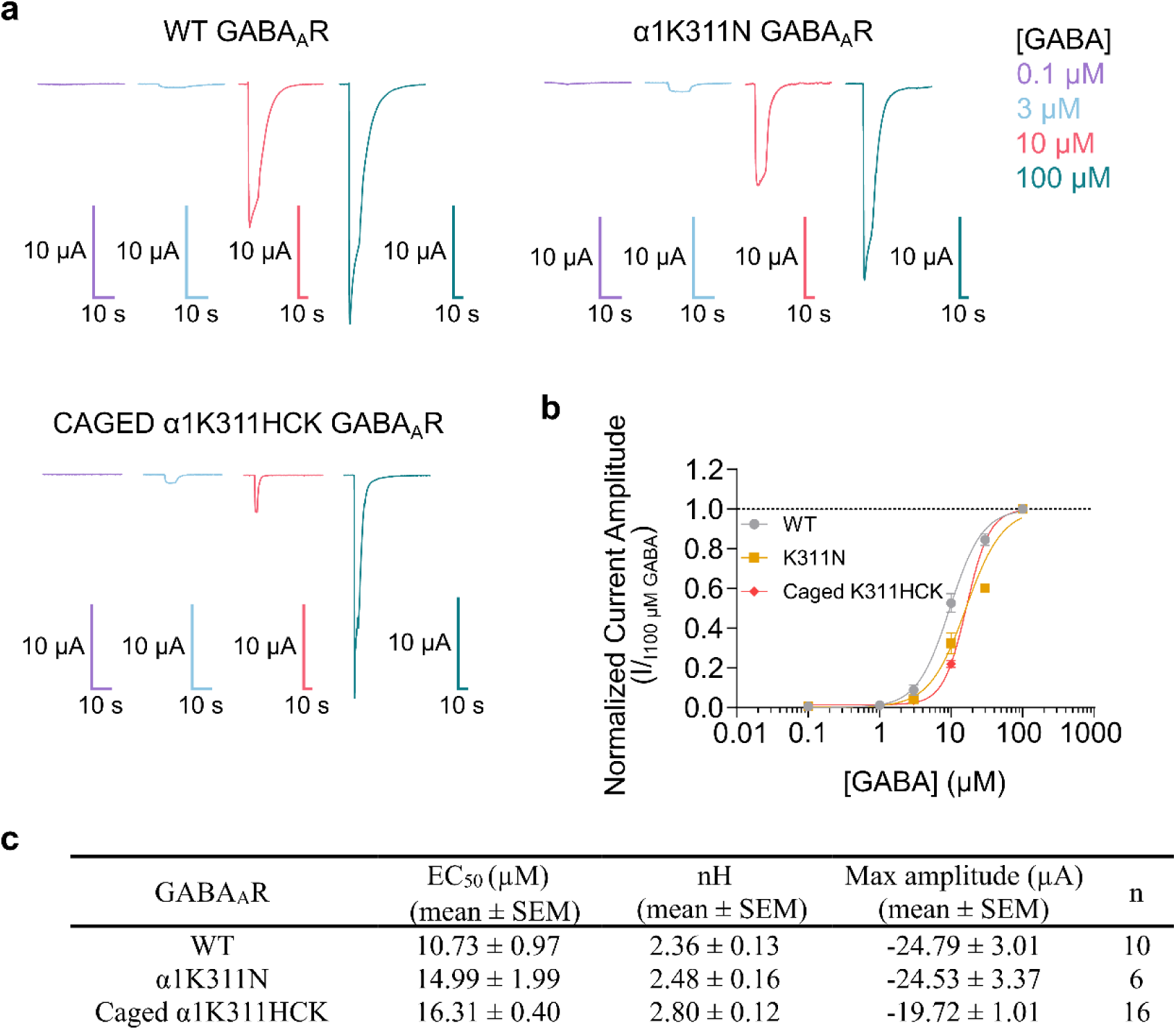
**Dose-response curve of GABA_A_R constructs.** (a) Representative traces correspond to current peak responses to varying concentrations (from 0.1 µM to 100 µM) of GABA. Top left, WT; top right, α1K311N; bottom left, Caged α1K311HCK. **(b)** Dose-response curves of WT, α1K311N, and caged α1K311HCK GABA_A_Rs were plotted. Data represents mean ± SEM. **(c)** Table shows EC_50_, Hill coefficient, max current amplitude, and sample information. Data were collected from n = 10–16 oocytes (4–5 batches).

**Extended Data Figure 6.**
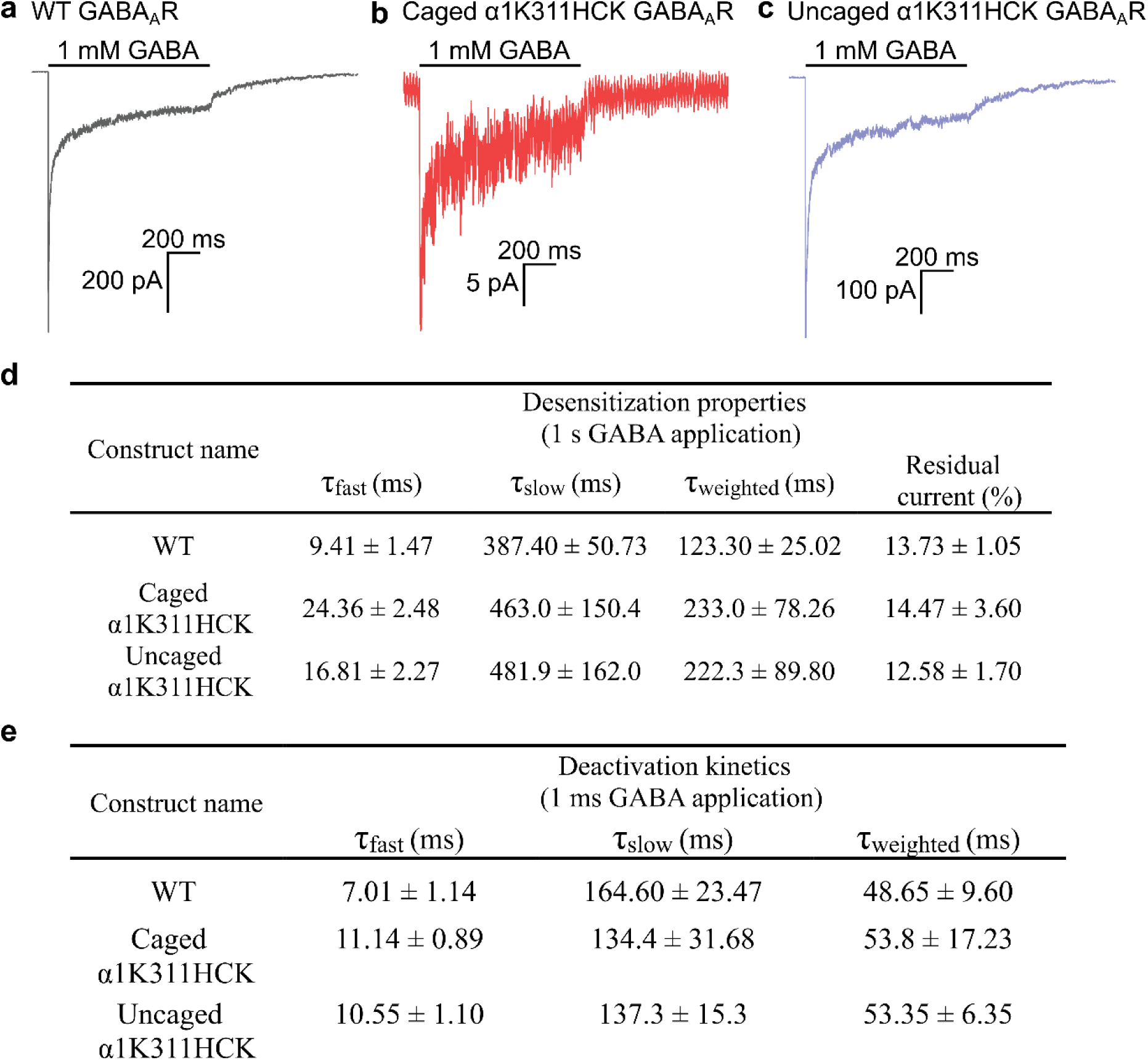
Outside-out patch-clamp recordings of WT and α1K311HCK GABA_A_Rs using rapid perfusion at physiological GABA concentration. **(a)** Outside-out patch-clamp recording traces in response to a 1-s application of 1 mM GABA with a 10-s interpulse interval for **(a)** WT, **(b)** caged α1K311HCK, and **(c)** uncaged α1K311HCK. Tables show desensitization **(d)** and deactivation **(e)** properties derived from double exponential fitted traces corresponding to (a–c) and Figure 4a,b, respectively. Data were collected from n = 6–8 patches (4 batches).

**Extended Data Figure 7.**
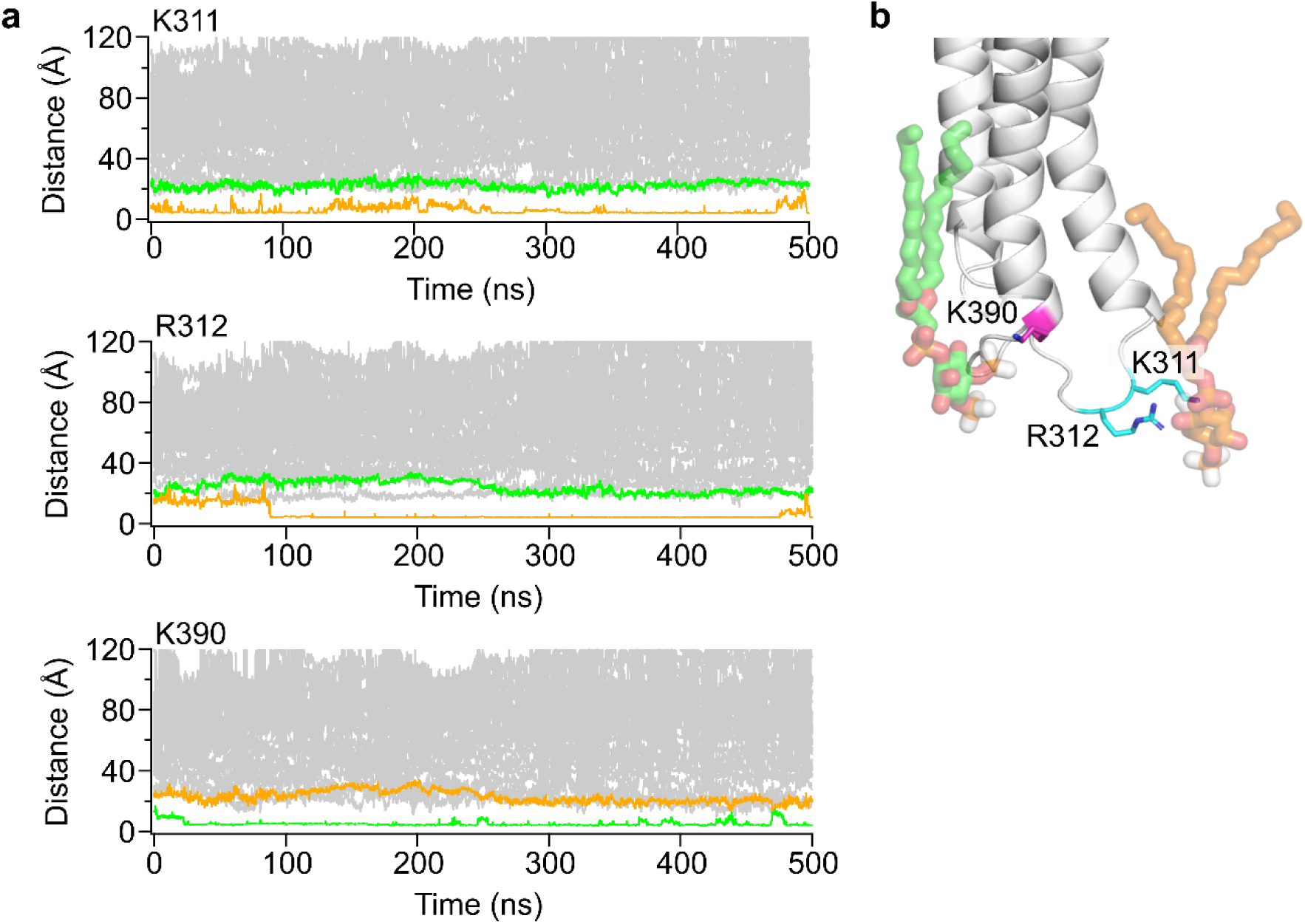
K390 interacts with PI(4,5)P_2_ molecules different from K311 and R312. **(a)** Distance measured from 5-phosphate of all PI(4,5)P_2_ molecules in the system to the terminal carbons of K311, R312, and K390. **(b)** Model of two separate PI(4,5)P_2_ molecules binding to K311/R312 (orange) and K390 (green).

**Extended Data Figure 8.**
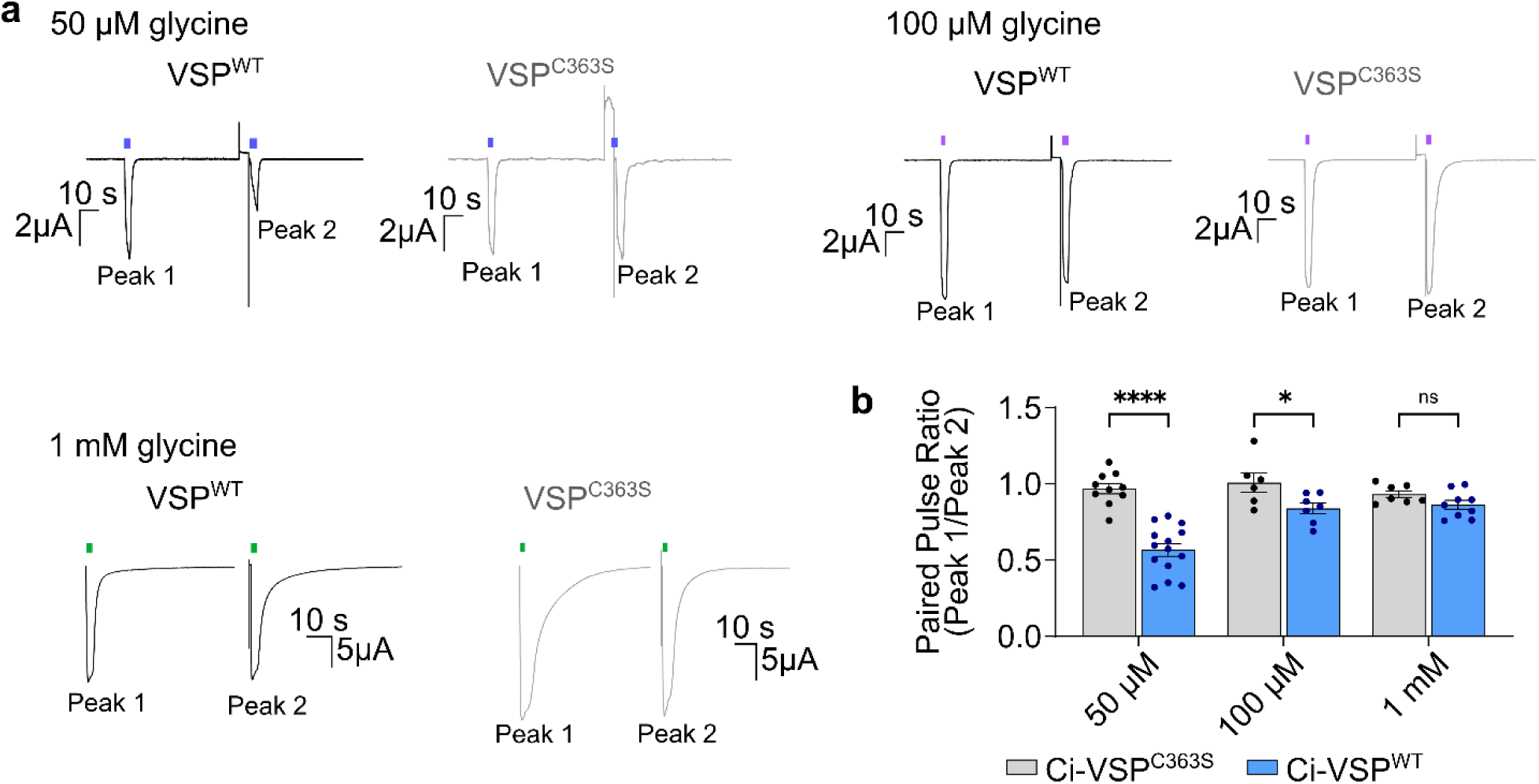
PI(4,5)P_2_ depletion reduces glycine-evoked responses of homopentameric α1 GlyR. **(a)** Representative traces of homopentameric α1 GlyR co-expressed with Ci-VSP; glycine-evoked currents with/without VSP activation at varying glycine concentrations (top, 50 μM glycine; middle, 100 μM glycine; bottom, 1 mM glycine). Glycine applications are indicated by bars. **(b)** Summary of paired pulse ratios compared between homopentameric α1 GlyR co-expressed with Ci-VSP and Ci-VSP^C363S^ mutant at different glycine concentrations. Statistical significance was assessed using ordinary two-way ANOVA followed by Bonferroni’s post hoc test (50 µM, p = <0.0001; 100 µM, p = 0.047; 1 mM, p = 0.798; n = 6–14 oocytes from 2–3 batches). Data represents mean ± SEM.

**Extended Data Figure 9.**
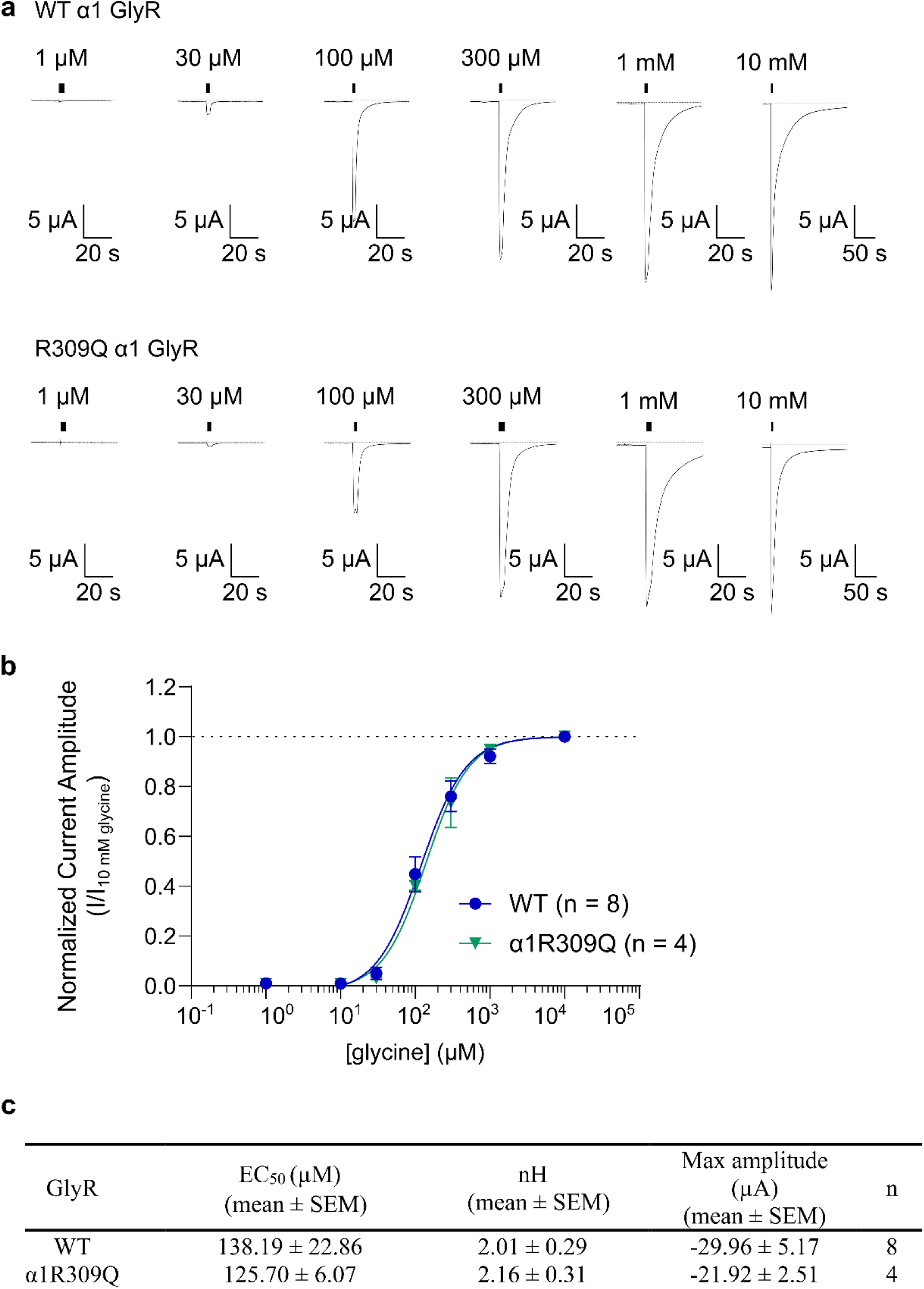
Homopentameric α1 GlyR (WT) vs α1R309Q mutant dose-response analysis. **(a)** Representative current responses to varying glycine concentrations. (Top) WT and (bottom) α1R309Q mutant. **(b)** Dose-response curves of WT and α1R309Q. Data represents mean ± SEM. **(c)** Table shows EC_50_, Hill coefficient, max current amplitude, and sample size information. Data were collected from n = 4–8 oocytes from 2–3 batches.

**Extended Data Figure 10.**
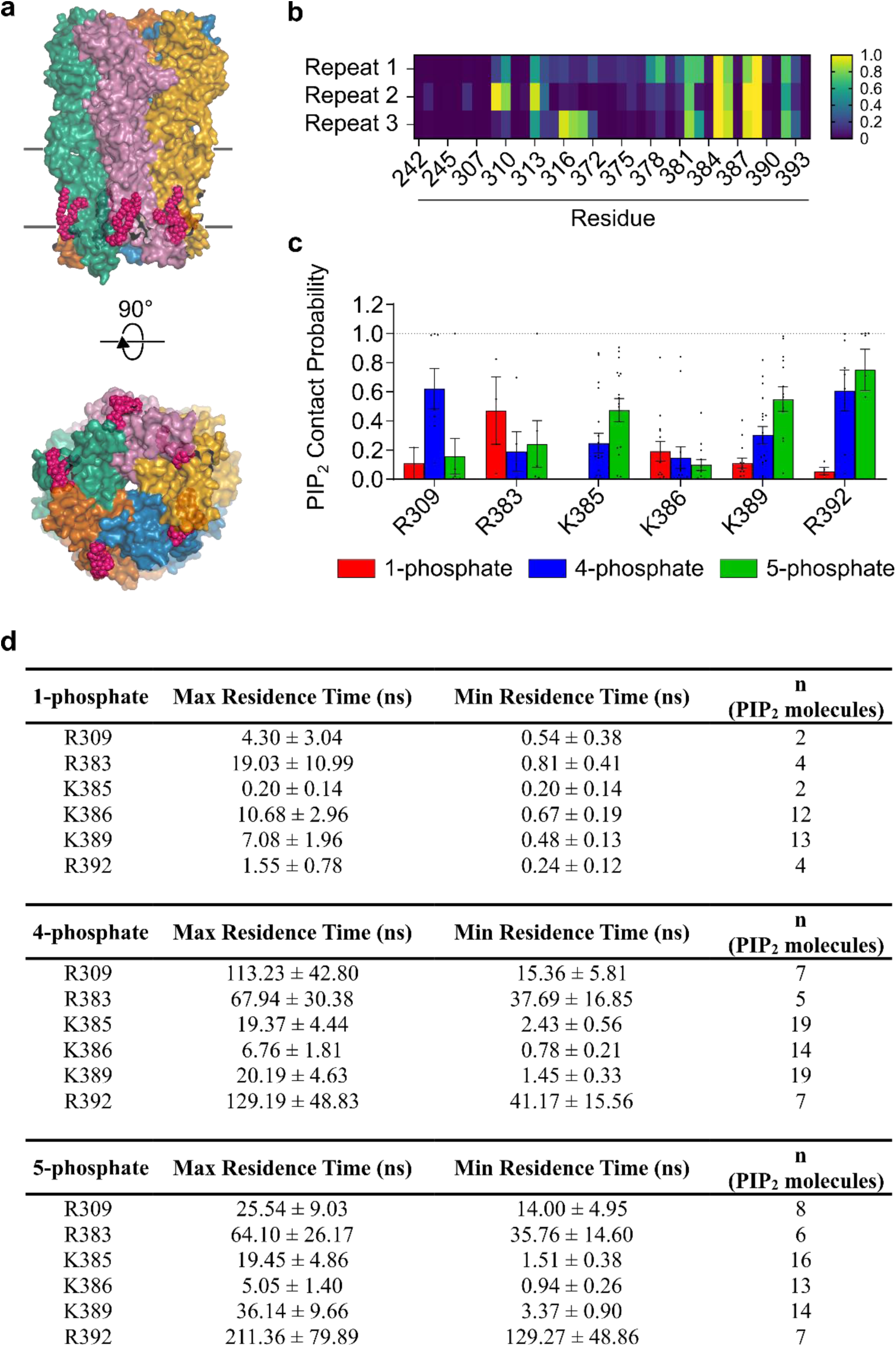
Homopentameric α1 GlyR bind PI(4,5)P_2_ in intersubunit sites. **(a)** Structural representative α1 GlyR shown in surface view. Subunits are colored individually and PI(4,5)P_2_ molecules are shown in stick (magenta). **(b)** Initial screening for PI(4,5)P_2_ contact site (distance ≤ 10 Å). **(c)** Summarized contact probability per putative binding sites to each phosphate. Contact was redefined as minimum distances ≤ 5 Å. Each data point represents an individual PI(4,5)P_2_ molecule. Data represents mean ± SEM. **(d)** Table shows max and mean residence times per residue (mean ± SEM).

