## Supplementary Information for "PI(4,5)P_2_-dependence of GABA_A_ receptor channel function revealed by optogenetic manipulation of a binding site"

**Supplementary Table 1. Comprehensive list of available GABA_A_R structures.**

List of GABA_A_R structure from PDB database. Structures that show PI(4,5)P_2_ as a one of the ligands are indicated with (*).

**Supplementary Table 2. Primer list for site-directed mutagenesis.**

List of forward and reverse primer sequences used to perform site-directed mutagenesis.

**Supplementary Table 3. PI(4,5)P_2_ contact probability, max and mean residence time (ns) information for R248, K311, R312, and K390 of WT GABA_A_R MD simulation.**

| 1-phosphate | PIP_2_ contact probability | Max Residence Time (ns) | Mean Residence Time (ns) | n  (PIP_2_ molecule) |
| --- | --- | --- | --- | --- |
| R248 | 0.11 ± NA | 4.70 ± NA | 1.15 ± NA | 1 |
| K311 | 0.04 ± 0.03 | 0.58 ± 0.40 | 0.17 ± 0.05 | 6 |
| R312 | 0.02 ± 0.01 | 1.53 ± 0.72 | 0.42 ± 0.17 | 3 |
| K390 | 0.13 ± 0.08 | 8.60 ± 4.26 | 0.71 ± 0.20 | 3 |
| 4-phosphate | PIP_2_ contact probability | Max Residence Time (ns) | Mean Residence Time (ns) | n  (PIP_2_ molecule) |
| R248 | 0.00 ± NA | 0.40 ± NA | 0.40 ± NA | 1 |
| K311 | 0.29 ± 0.09 | 9.62 ± 3.87 | 0.54 ± 0.11 | 9 |
| R312 | 0.04 ± 0.02 | 1.67 ± 0.64 | 0.31 ± 0.13 | 6 |
| K390 | 0.20 ± 0.13 | 9.93 ± 8.49 | 1.00 ± 0.71 | 7 |
| 5-phosphate | PIP_2_ contact probability | Max Residence Time (ns) | Mean Residence Time (ns) | n  (PIP_2_ molecule) |
| R248 | NA | NA | NA | NA |
| K311 | 0.45 ± 0.12 | 44.83 ± 19.42 | 7.33 ± 4.54 | 9 |
| R312 | 0.72 ± 0.12 | 113.08 ± 12.73 | 19.40 ± 3.07 | 6 |
| K390 | 0.51 ± 0.17 | 25.88 ± 12.02 | 1.78 ± 0.36 | 6 |

Supplementary Table 3 provides averaged values (across three replicates) ± SEM of PI(4,5)P_2_ contact probability, max and mean residence time (ns) in WT GABA_A_R simulations.

**Supplementary Table 4. PI(4,5)P_2_ contact probability, max and mean residence time (ns) information for R248, K311, R312, and K390 of α1K311N GABA_A_R MD simulation.**

| 1-phosphate | PIP_2_ contact probability | Max Residence Time (ns) | Mean Residence Time (ns) | n  (PIP_2_ molecule) |
| --- | --- | --- | --- | --- |
| R248 | NA | NA | NA | NA |
| N311 | 0.29 ± NA | 3.20 ± NA | 0.28 ± NA | 1 |
| R312 | 0.01 ± 0.01 | 0.30 ± 0.10 | 0.15 ± 0.05 | 2 |
| K390 | 0.12 ± 0.08 | 5.92 ± 3.73 | 0.54 ± 0.14 | 5 |
| 4-phosphate | PIP_2_ contact probability | Max Residence Time (ns) | Mean Residence Time (ns) | n  (PIP_2_ molecule) |
| R248 | 0.24 ± 0.12 | 22.67 ± 11.29 | 7.60 ± 5.94 | 3 |
| N311 | 0.03 ± 0.02 | 1.02 ± 0.29 | 0.22 ± 0.03 | 6 |
| R312 | 0.13 ± 0.07 | 4.58 ± 2.98 | 0.27 ± 0.08 | 4 |
| K390 | 0.12 ± 0.05 | 9.24 ± 6.32 | 0.66 ± 0.33 | 11 |
| 5-phosphate | PIP_2_ contact probability | Max Residence Time (ns) | Mean Residence Time (ns) | n  (PIP_2_ molecule) |
| R248 | 0.83 ± 0.17 | 218.15 ± 66.85 | 42.20 ± 37.68 | 2 |
| N311 | 0.03 ± 0.03 | 0.50 ± 0.40 | 0.15 ± 0.05 | 3 |
| R312 | 0.61 ± 0.14 | 75.64 ± 22.32 | 14.81 ± 6.02 | 5 |
| K390 | 0.40 ± 0.08 | 35.86 ± 13.14 | 4.13 ± 1.74 | 8 |

Supplementary Table 4 provides averaged values (across three replicates) ± SEM of PI(4,5)P_2_ contact probability, max and mean residence time (ns) in α1K311N GABA_A_R simulations.

**Supplementary Table 5. Comprehensive list of available GlyR structures.**

List of GlyR structures from PDB database.

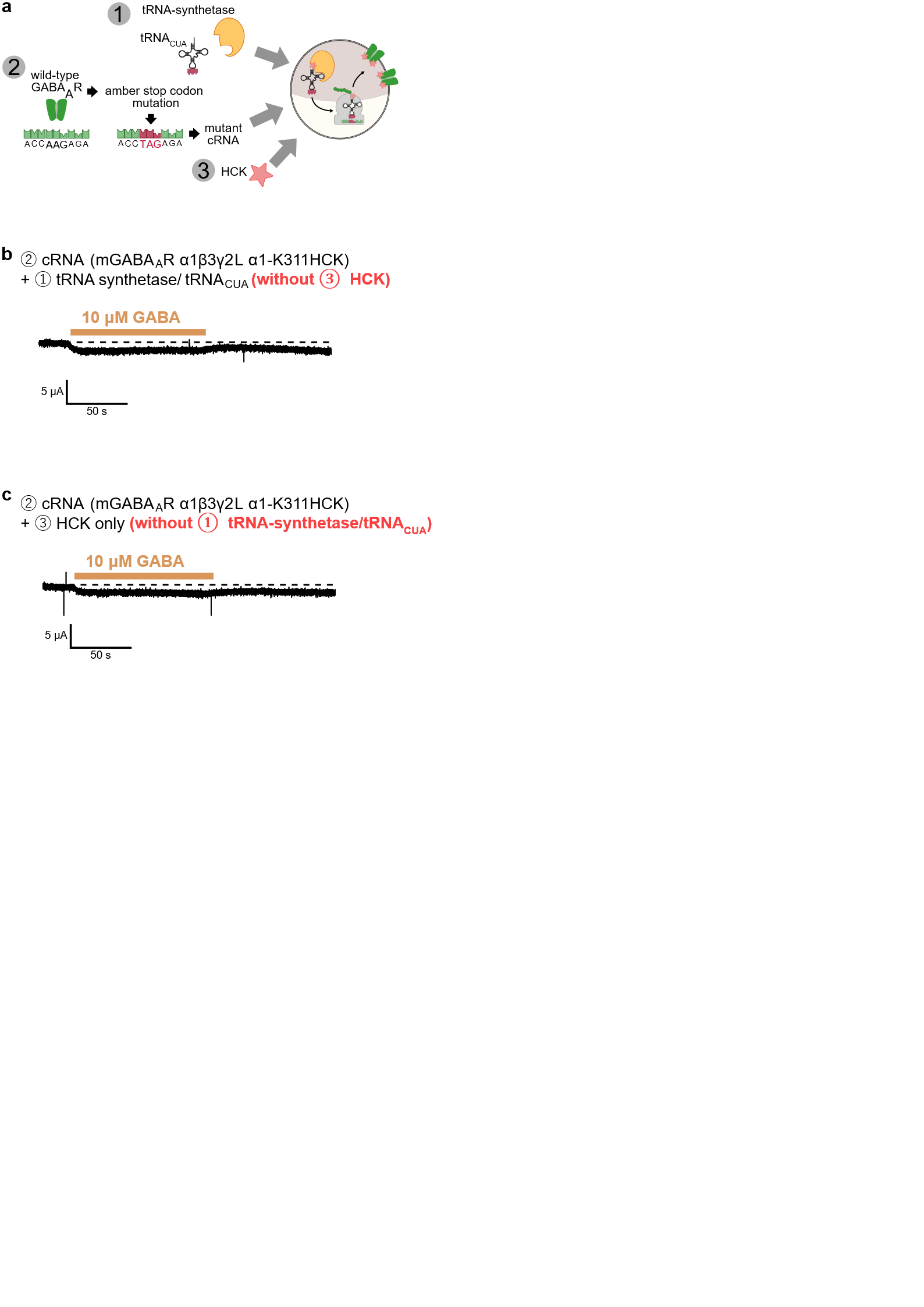

**Supplementary Figure 1. Omitting either the HCK or tRNA confirms the current and the current increase during uncaging of α1K311HCK is solely from the successful incorporation of caged lysine.**

**(a)** Schematic overview of HCK incorporation in *Xenopus* oocytes. Omitting components **(b)** HCK or **(c)** tRNA-synthetase and tRNA_CUA_ result in negligible GABA-evoked current.

**SUPPLEMENTARY INFORMATION REFERENCES**
